# Automated long-term recording and analysis of neural activity in behaving animals

**DOI:** 10.1101/033266

**Authors:** Ashesh K. Dhawale, Rajesh Poddar, Evi Kopelowitz, Valentin Normand, Steffen B. E. Wolff, Bence P. Ölveczky

**Affiliations:** Department of Organismic and Evolutionary Biology and Center for Brain Science, Harvard University, Cambridge MA, USA 02138

**Author notes:** co-first authors (Ashesh K. Dhawale, Rajesh Poddar).

## Abstract

Addressing how neural circuits underlie behavior is routinely done by measuring electrical activity from single neurons during experimental sessions. While such recordings yield snapshots of neural dynamics during specified tasks, they are ill-suited for tracking single-unit activity over longer timescales relevant for most developmental and learning processes, or for capturing neural dynamics across different behavioral states. Here we describe an automated platform for continuous long-term recordings of neural activity and behavior in freely moving animals. An unsupervised algorithm identifies and tracks the activity of single units over weeks of recording, dramatically simplifying the analysis of large datasets. Months-long recordings from motor cortex and striatum made and analyzed with our system revealed remarkable stability in basic neuronal properties, such as firing rates and inter-spike interval distributions. Interneuronal correlations and the representation of different movements and behaviors were similarly stable. This establishes the feasibility of high-throughput long-term extracellular recordings in behaving animals.

**Highlights:**

- We record neural activity and behavior in rodents continuously (24/7) over months
- An automated spike-sorting method isolates and tracks single units over many weeks
- Neural dynamics and motor representations are highly stable over long timescales
- Neurons cluster into functional groups based on their activity in different states

**eTOC Blurb:** Dhawale et al. describe experimental infrastructure for recording neural activity and behavior continuously over months in freely moving rodents. A fully automated spike-sorting algorithm allows single units to be tracked over weeks of recording. Recordings from motor cortex and striatum revealed a remarkable long-term stability in both single unit activity and network dynamics.

## Introduction

The goal of systems neuroscience is to understand how neural activity generates behavior. A common approach is to record from neuronal populations in targeted brain areas during experimental sessions while subjects perform designated tasks. Such intermittent recordings provide brief ‘snapshots’ of task-related neural dynamics (Georgopoulos et al., 1986; Hanks et al., 2015; Murakami et al., 2014), but fail to address how neural activity is modulated outside of task context and across behavioral states (Arieli et al., 1996; Evarts, 1964; Gomez-Marin et al., 2014; Gulati et al., 2014; Lee and Dan, 2012; Rinberg et al., 2006; Santhanam et al., 2007; Wilson and McNaughton, 1994). Furthermore, intermittent recordings are ill-suited for reliably tracking the same neurons over time (Dickey et al., 2009; Fraser and Schwartz, 2012; McMahon et al., 2014; Santhanam et al., 2007; Tolias et al., 2007), making it difficult to discern how neural dynamics and properties of individual neurons are shaped by developmental and learning processes that evolve over longer timescales (Ganguly et al., 2011; Jog et al., 1999; Lütcke et al., 2013; Marder and Goaillard, 2006; Peters et al., 2014; Singer et al., 2013).

Addressing such fundamental questions would be greatly helped by recording neural activity and behavior continuously over days and weeks in freely moving animals. Such longitudinal recordings would allow us to follow the activity of single neurons over more trials, experimental conditions, and behavioral states, thus increasing the power with which inferences about neural function can be made (Lütcke et al., 2013). While *in vivo* calcium imaging allows the same neuronal population to be recorded intermittently over long durations (Huber et al., 2012; Peters et al., 2014; Ziv et al., 2013), photobleaching, phototoxicity, and cytotoxicity (Grienberger and Konnerth, 2012; Looger and Griesbeck, 2012), as well as the requirements for head-restraint in many versions of such experiments (Dombeck et al., 2007; Huber et al., 2012; Peters et al., 2014), make the method unsuitable for continuous long-term recordings. Calcium imaging also has relatively poor temporal resolution (Grienberger and Konnerth, 2012; Vogelstein et al., 2009), limiting its ability to resolve precise spike patterns (Vogelstein et al., 2010; Yaksi and Friedrich, 2006) (but see Gong et al., 2015 for an alternative high-speed voltage sensor). In contrast, extracellular recordings using electrode arrays can measure the activity of many single neurons simultaneously with sub-millisecond resolution (Buzsáki, 2004). Despite the unique benefits of continuous (24/7) long-term electrical recordings, they are not routinely performed. A major reason is the inherently laborious and difficult process of reliably and efficiently tracking the activity of single units from such longitudinal datasets (Einevoll et al., 2012), wherein firing rates of individual neurons can vary over many orders of magnitude (Hromádka et al., 2008; Mizuseki and Buzsáki, 2013) and spike waveforms change over time (Dickey et al., 2009; Emondi et al., 2004; Fraser and Schwartz, 2012; Okun et al., 2015a; Santhanam et al., 2007; Tolias et al., 2007).

To address this, we designed and deployed a low-cost recording system that enables fully automated long-term continuous extracellular recordings from large numbers of neurons in freely behaving rodents engaged in natural behaviors and prescribed tasks. To efficiently parse the large streams of neural data, we developed an unsupervised spike sorting algorithm that automatically processes the raw data from electrode array recordings, and clusters spiking events into putative single units, tracking their activity over long timescales. We used this integrated system to record from motor cortex and striatum continuously over several months. These experiments revealed a remarkable stability in basic neuronal properties, such as firing rates and inter-spike interval distributions. Interneuronal correlations and movement tuning across a range of behaviors were similarly stable. We also found that neurons within a brain region appear to be organized into distinct functional groups based on their activity patterns across behavioral states and time.

## Results

### Infrastructure for automated long-term neural recordings in behaving animals

We developed experimental infrastructure for continuous long-term extracellular recordings in behaving rodents. Our starting point was ARTS, a fully Automated Rodent Training System we previously developed (Poddar et al., 2013). In ARTS, the animal’s home-cage doubles as the experimental chamber, making it a suitable platform for continuous long-term recordings.

To ensure that animals remain reliably and comfortably connected to the recording apparatus over months-long experiments, we designed a tethering system that allows experimental subjects to move around freely while preventing them from reaching for (and chewing) the signal cable (Figure 1A; see Supplemental Experimental Procedures for details). Our solution connects the implanted recording device via a cable to a passive commutator attached to a carriage that rides on a low-friction linear slide. The carriage is counterweighted by a pulley, resulting in a small constant upwards force (< 10 g) on the cable that keeps it taut and out of the animal’s reach without unduly affecting its movements. The recording extension can easily be added to our custom home-cages, allowing animals that have been trained, prescreened, and deemed suitable for recordings to be implanted with electrode drives, and placed back into their familiar training environment (i.e. home-cage) for recordings.

**Figure 1:**
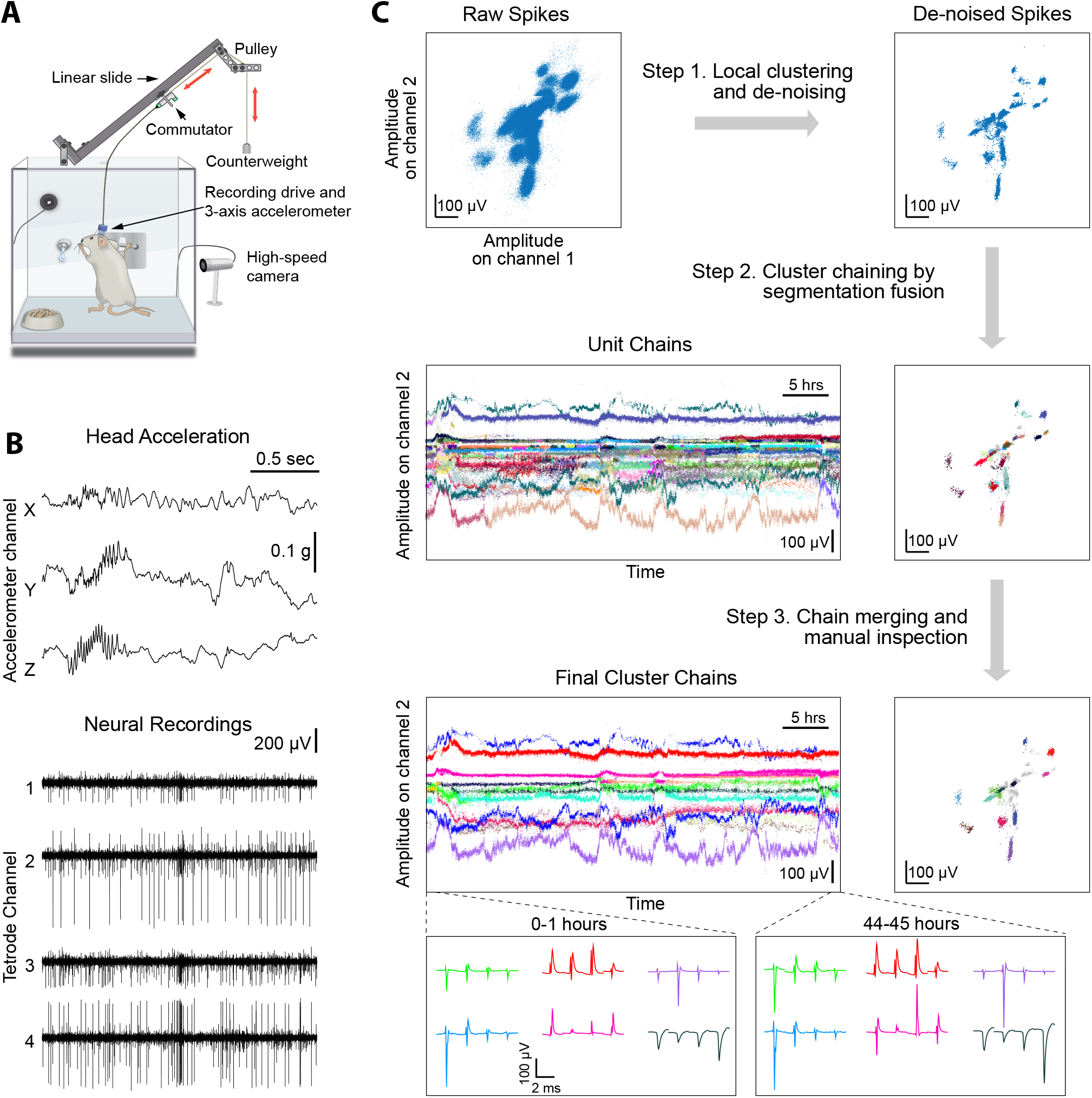
Experimental infrastructure and analysis pipeline for long-term continuous neural and behavioral recordings in behaving rodents. A. Adapting our automated rodent training system (ARTS) for long-term electrophysiology. Rats engage in natural behaviors and prescribed motor tasks in their home-cages, while neural data is continuously acquired from implanted electrodes. The tethering cable connects the head-stage to a commutator mounted on a carriage that moves along a low-friction linear slide. The commutator-carriage is counterweighted to eliminate slack in the tethering cable. Behavior is continuously monitored and recorded using a camera and a 3-axis accelerometer.
B. Example of a recording segment showing high-resolution behavioral and neural data simultaneously acquired from a head-mounted 3-axis accelerometer and a tetrode implanted in the motor cortex, respectively.
C. Overview of our fully automated spike tracker (FAST) (see Experimental Procedures). (Top, Left) Spike amplitudes on two electrodes of an example tetrode for a 1 hour-long recording segment. Clusters of spikes with similar waveforms are identified and tracked over time. An iterative local-clustering step (Step 1) compresses and de-noises the dataset. De-noised spike clusters (Top, right) are linked across time by a segmentation fusion algorithm (Step 2) to yield cluster-chains corresponding to single units (middle). In the final step, we visually inspect the output of the automated sorting and merge similar chains across time (Step 3) to yield the final single-unit clusters (bottom). Insets (bottom) show average spike waveforms of six example units 44 hours apart.

Extracellular signals recorded from behaving animals are amplified and digitized on a custom-designed head-stage (Figure 1B, Supplemental Experimental Procedures). To characterize the behavior of animals throughout the recordings, the head-stage features a 3- axis accelerometer that measures head movements at high temporal resolution (Venkatraman et al., 2010) (Figure 1A-B). We also record continuous video of the rats’ behavior with a wideangle camera above the cage (Experimental Procedures). The large volumes of behavioral and neural data (∼0.5 TB/day/rat) are streamed to custom-built high-capacity servers.

### A Fully Automated Spike Tracker (FAST) for long-term neural recordings

Extracting single-unit spiking activity from raw data collected over weeks and months of continuous extracellular recordings presents a significant challenge for which there is currently no adequate solution. Parsing such large datasets must necessarily rely on automated spikesorting methods. These face three major challenges (Rey et al., 2015): First, they must reliably capture the activity of simultaneously recorded neurons whose firing rates can vary over many orders of magnitude (Hromádka et al., 2008; Mizuseki and Buzsáki, 2013). Second, they have to contend with spike shapes from recorded units changing significantly over time (Dickey et al., 2009; Emondi et al., 2004; Fraser and Schwartz, 2012). Third, they must be capable of processing very large numbers of spikes in a reliable and efficient manner (in our experience >10^10^ spikes per rat over a time span of 3 months for 64 channel recordings).

Here we present an unsupervised spike sorting algorithm that meets these challenges and allows single units to be tracked over months-long timescales. Our Fully Automated Spike Tracker (FAST) comprises two main steps. First, to compress the datasets and normalize for large variations in firing rates between units, it applies ‘local clustering’ to create denoised representations of spiking events in the data. In a second step, FAST chains together de-noised spikes belonging to the same putative single unit over time using an integer linear programming algorithm (Vazquez-Reina et al., 2011). FAST is an efficient and modular data processing pipeline that, depending on overall spike rates, runs two to three times faster than the rate at which the data (64 electrodes) is acquired.

To parse and compress the raw data, FAST first identifies and extracts spike events (‘snippets’) by bandpass filtering and thresholding each electrode channel (Supplemental Experimental Procedures, Figure S1). Spike clustering is then performed on blocks of 1000 consecutive spike ‘snippets’ by means of an automated superparamagnetic clustering routine, a step we call ‘local clustering’ (Blatt et al., 1996; Quiroga et al., 2004) (Supplemental Experimental Procedures, Figure 1C and S2). Spikes in a block that belong to the same cluster are replaced by their centroid, a step that effectively de-noises and compresses the data by representing groups of similar spikes with a single waveform. However, due to large differences in firing rates across different units, the initial blocks of 1000 spikes will be dominated by high firing rate units. Spikes from more sparsely firing cells that do not contribute at least 15 spikes to a cluster in a given block are carried forward to the next round of local clustering, where previously assigned spikes have been removed (Supplemental Experimental Procedures, Figure S2). Applying this method of pooling and local clustering sequentially four times generates a denoised dataset that accounts for large differences in the firing rates of simultaneously recorded units (Figure 1C, Supplemental Experimental Procedures).

The second step of the FAST algorithm is inspired by an automated method (‘segmentation fusion’) that links similar elements over cross-sections of longitudinal datasets in a globally optimal manner (Supplemental Experimental Procedures, Figure S3). Segmentation fusion has been used to trace processes of individual neurons across stacks of two-dimensional serial electron-microscope images (Kasthuri et al., 2015; Vazquez-Reina et al., 2011). We adapted this method to link similar de-noised spikes across consecutive blocks into *chains* containing the spikes of putative single units over longer timescales (Figure 1C). This algorithm allows us to automatically track the same units over days and weeks of recording.

In a final post-processing and verification step, we use a semi-automated method to link ‘chains’ belonging to the same putative single unit together, and perform visual inspection of each unit. A detailed description of the various steps involved in the automated spike sorting can be found in Supplemental Experimental Procedures. Below, we describe how our algorithm parses data acquired from continuous long-term tetrode recordings, but we note that it can be adapted to efficiently analyze other types of electrode array or single electrode recordings – whether continuous or intermittent.

### Continuous long-term recordings from striatum and motor cortex

To demonstrate the utility of our experimental platform and analysis pipeline for long-term neural recordings, we implanted tetrode drives (16 tetrodes, 64 channels) into dorsolateral striatum (n=1) or motor cortex (n=1) of rats (Experimental Procedures). We recorded electrophysiological and behavioral data continuously, or with only brief interruptions, for more than 3 months. We note that our recordings terminated not because of technical issues with the implant or recording apparatus, but because we deemed the experiments to have reached their end points.

We used our automated spike sorting method (FAST) to cluster spike waveforms into putative single units, isolating a total of 1031 units from motor cortex and 719 units from striatum (Figure 2A). On average, we could track single units over several days (mean: 3.5 days for motor cortex and 4.4 days for striatum), with significant fractions of units tracked continuously for more than a week (18% and 20% in motor cortex and striatum, respectively) and even a month (0.4% in motor cortex, Figure 2B). Periods of stable recordings were interrupted by either intentional advancement of the electrode drive or spontaneous ‘events’ likely related to the sudden movement of the electrodes (Figure 2A). On average, we recorded simultaneously from 30.2 ± 15.2 units in motor cortex and 27.2 ± 18.9 units in striatum (mean ± standard deviation) (Figure 2C).

**Figure 2:**
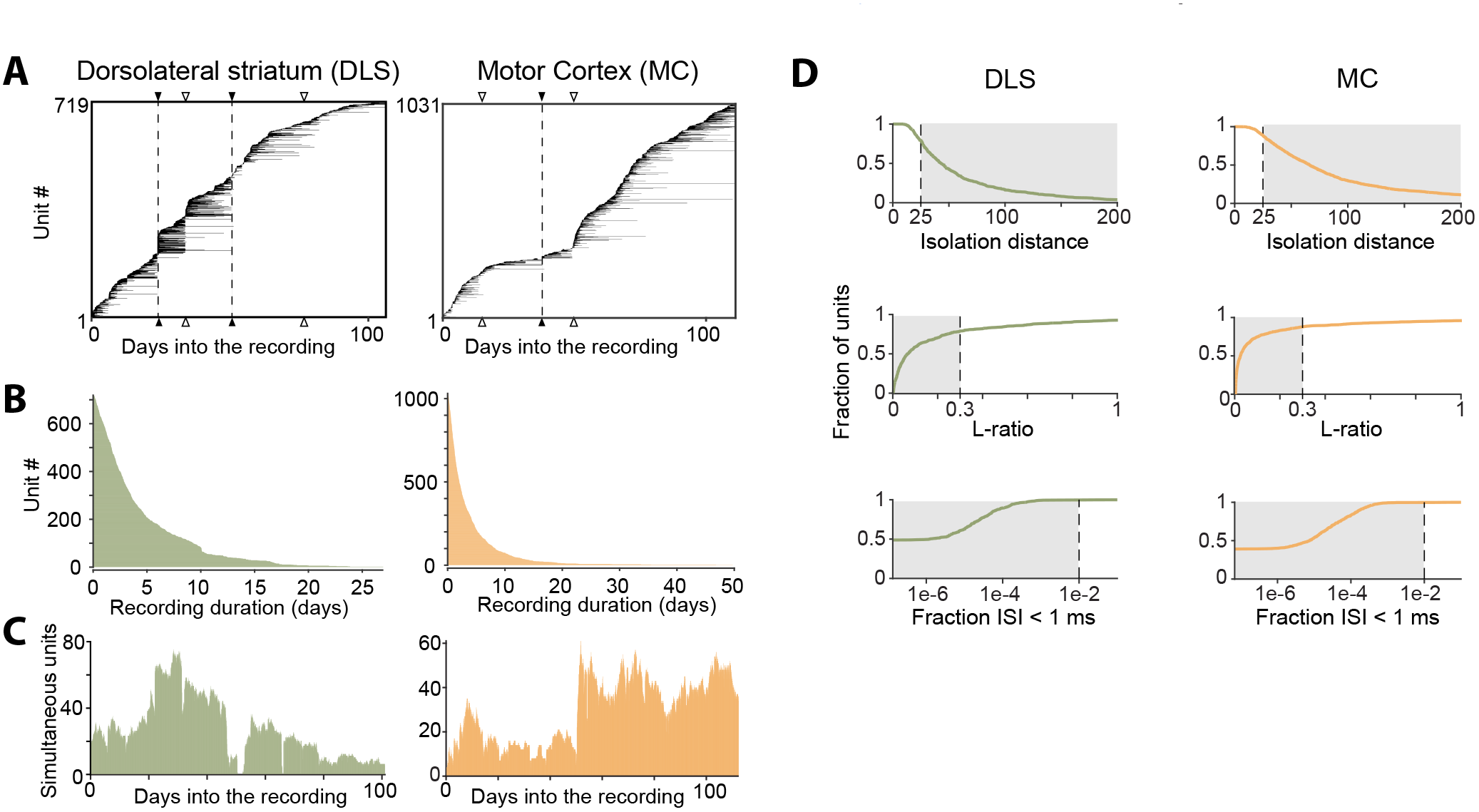
Single units isolated and tracked from continuous months-long recordings made in dorsolateral striatum (DLS) and motor cortex (MC) of behaving rats. A. Temporal profile of units recorded in DLS (left) and MC (right) over a period of ∼3 months. Unit recording times are indicated by black bars, and are sorted by when they were first identified in the recording. Black triangles and dotted lines indicate times at which the electrode array was intentionally advanced into the brain by turning the micro-drive. Open triangles indicate times at which the population of recorded units changed spontaneously.
B. Holding times for units recorded in the DLS (left, green) and MC (right, orange), sorted by duration.
C. Number of simultaneously recorded units in the DLS (left, green) and MC (right, orange) as a function of time in the recording.
D. Cumulative distributions of average cluster isolation quality for all units recorded in DLS (left, green) and MC (right, orange). Cluster quality was measured by the isolation distance (top), L-ratio (middle), and fraction of inter-spike intervals under 1 ms (bottom). Dotted lines mark the quality thresholds for each of these measures. Shaded regions denote acceptable values.

The quality of single unit isolation was assessed by computing quantitative measures of cluster quality, i.e. cluster isolation distance (Harris et al., 2000), L-ratio (Schmitzer-Torbert et al., 2005), and presence of a clean refractory period (Hill et al., 2011; Lewicki, 1998) (Figure 2D). Assessing the mean cluster quality of the units over their recording lifetimes, we found that 87.0% of motor cortical (n=897) and 77.5% of striatal units (n=554) satisfied our conservative criteria (Quirk et al., 2009; Schmitzer-Torbert et al., 2005; Sigurdsson et al., 2010) (Isolation distance >= 25, L-ratio <= 0.3 and fraction of ISIs below 1 ms <= 1%). However, 98.1% of motor cortical units (n=1011) and 96.2% of striatal units (n=692) met these criteria for at least one hour of recording.

Absent any ground truth information with which to benchmark the performance of our automated algorithm (Rey et al., 2015), we compared clusters identified by the FAST algorithm to those obtained from state-of-the art manual spike sorting within specified time-windows (Experimental Procedures). FAST successfully identified 72.1% of manually sorted spike clusters that exceeded our cluster quality criteria (n = 31 of 43 total), and 90.0 % of higher-quality clusters (Isolation distance >= 35, L-ratio <= 0.1, fraction of ISIs below 1 ms <= 1%; n = 18 of 20 total, Figure S4A). For clusters that were matched across the different sorting methods (identified by having more than 50% spike overlap), FAST was able to recover 93.3 ± 4.4% (median ± median absolute deviation) of spikes identified by manual sorting, and 95.5 ± 2.2% for high-quality clusters (Figure S4B). Finally, the isolation quality of matched clusters was, on average, similar for FAST and manual clustering (Figure S4C), suggesting that FAST is comparable to manual sorting in terms of cluster quality.

Previous attempts to track populations of extracellularly recorded units over time relied on matching units across discontinuous recording sessions based on similarity metrics such as spike waveform distance (Dickey et al., 2009; Emondi et al., 2004; Fraser and Schwartz, 2012; Ganguly and Carmena, 2009; Greenberg and Wilson, 2004; Jackson and Fetz, 2007; McMahon et al., 2014; Okun et al., 2015a; Thompson and Best, 1990; Tolias et al., 2007) and activity measures including firing rates (Fraser and Schwartz, 2012) and inter-spike interval histograms (Dickey et al., 2009). Given that spike waveforms of single-units can undergo significant changes even within a day (Figure 1C), a major concern with discontinuous tracking methods is the difficulty in verifying their performance. Since we were able to reliably track units continuously over weeks, we used the sorted single-units as ‘ground truth’ data with which to benchmark the performance of discontinuous tracking methods (Figure S4D-E). In our data set, we found discontinuous tracking to be highly error-prone: applying tolerant thresholds for merging units across time, a large fraction of distinct units were labeled as the same (false positives), while at conservative thresholds only a small proportion of the units were reliably tracked across days (Figure S4E).

The ability to record and analyze the activity of large populations of single units continuously over days and weeks allows neural processes that occur over long time-scales to be interrogated. Below we analyze and describe single neuron activity, interneuronal correlations, and the relationship between neuronal dynamics and different behavioral states, over weeks-long time scales.

### Stability of neural dynamics

We clustered motor cortical and striatal units into putative principal neurons and interneurons based on their spike shapes and firing rates (Barthó et al., 2004; Berke et al., 2004; Connors and Gutnick, 1990), thus identifying 264 fast spiking and 455 medium spiny neurons in striatum, and 591 fast spiking and 440 regular spiking neurons in motor cortex (Figure 3A). Consistent with previous reports (Hromádka et al., 2008; Mizuseki and Buzsáki, 2013), average firing rates were log-normally distributed and varied over more than three orders of magnitude (from 0.029 Hz to 20.5 Hz in striatum, and 0.031 Hz to 37.2 Hz in motor cortex; Figure 3A).

**Figure 3:**
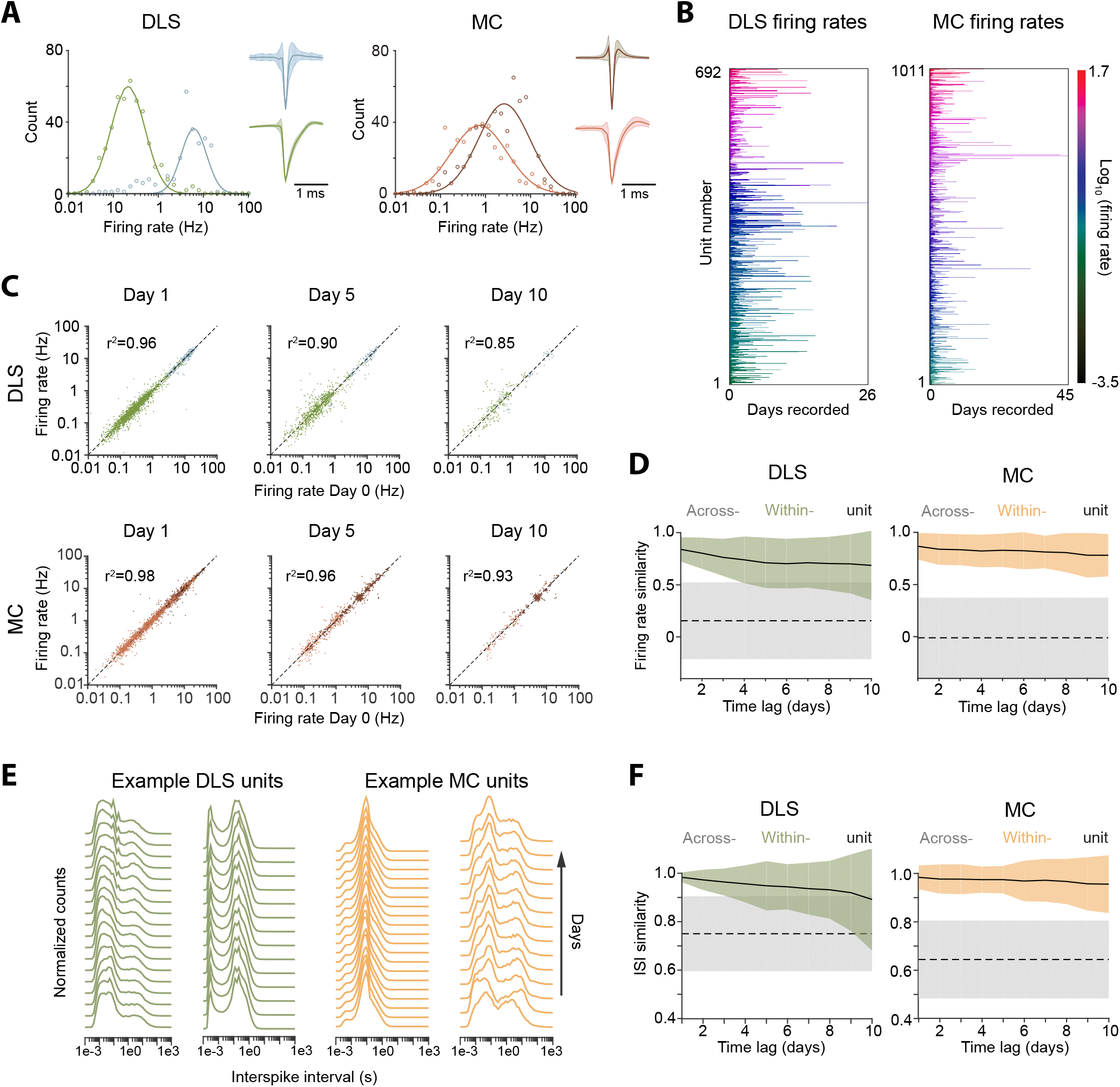
Long-term stability of single unit activity A. Histograms of average firing rates for units recorded in DLS (left) and MC (right). Putative cell-types, medium spiny neurons (MSN, blue) and fast-spiking interneurons (FSI, green) in DLS, and regular spiking (RS, brown) and fast spiking (FS, red) neurons in the MC, were classified based on spike shape and firing rate (Supplemental Experimental Procedures). The continuous traces are log-normal fits to the firing rate distributions of each putative cell-type. Insets show average peak-normalized waveform shapes for MSNs (left-bottom) and FSIs (left-top), and RS (right-bottom) and FS (right-top) neurons. Shading represents the standard deviation of the spike waveforms.
B. Firing rates of DLS (left) and MC (right) units for their recording duration. The color scale indicates firing rate on a log-scale, calculated in one-hour blocks. Units have been sorted by average firing rate.
C. Scatter plots of unit firing rates over time-lags of 1 (left), 5 (middle) and 10 (right) days for DLS (top) and MC (bottom). The dashed lines indicate equality. Every dot is a comparison of a unit’s firing from a baseline day to 1, 5 or 10 days later. The color of the dot indicates putative cell-type as per (A). Each unit may contribute multiple data points, depending on the length of the recording. Day 1: n = 2024 comparisons for striatum and n = 2225 for cortex; Day 5: n = 851 comparisons for striatum and n = 897 for cortex; Day 10: n = 255 comparisons for striatum and n = 350 for cortex.
D. Stability of unit firing rates over time. The firing rate similarity (see Supplemental Experimental Procedures) was measured across time-lags of 1 to 10 days for the same unit (within-unit, solid lines), or between simultaneously recorded units (across-unit, dashed lines) in DLS (left) and MC (right). Colored shaded regions indicate the standard deviation of within-unit firing rate similarity, over all units. Grey shaded regions indicate standard deviation of across-unit firing rate similarity, over all time-bins.
E. Inter-spike interval (ISI) histograms for example units in DLS (left, green) and MC (right, orange) over two weeks of continuous recordings. Each line represents the normalized ISI histogram measured on a particular day.
F. Stability of unit ISI distributions over time. Correlations between ISI distributions were measured across time-lags of 1 to 10 days for the same unit (within-unit, solid lines), or between simultaneously recorded units (across-unit, dashed lines) in MC (left) and DLS (right). Colored shaded regions indicate the standard deviation of within-unit ISI similarity, over all units. Grey shaded regions indicate standard deviation of across-unit ISI similarity, over all time-bins.

Average firing rates remained largely unchanged even over 10 days of recording (Figure 3B-D), suggesting that activity levels of single units in both cortex and striatum are stable over long timescales, and that individual units maintain their own firing rate set-point (Hengen et al., 2013; Marder and Goaillard, 2006). We next asked whether neurons also maintain second-order spiking statistics. The inter-spike interval (ISI) distribution is a popular metric that is sensitive to a cell’s mode of spiking (bursting, tonic etc.) and other intrinsic properties, such as refractoriness and spike frequency adaptation. Similarly to firing rate, we found that the ISI distribution of single units remained largely unchanged across days (Figure 3E-F).

Measures of single unit activity do not adequately address the stability of the network in which the neurons are embedded, as they do not account for interneuronal correlations (Abbott and Dayan, 1999; Ecker et al., 2010; Nienborg et al., 2012; Okun et al., 2015b; Salinas and Sejnowski, 2001; Singer, 1999). To address this, we calculated the cross-correlograms of all simultaneously recorded neuron pairs (Figure 4A, Supplemental Experimental Procedures). We found that 29.9% of striatal pairs (n=21990 pairs) and 46.7% motor cortex pairs (n=32567 pairs) were significantly correlated (Supplemental Experimental Procedures). The average time-lag of significantly correlated pairs was 26.3 ± 43.1 ms (median: 8.0 ms) and 36.6 ± 47.9 ms (median: 16.0 ms) for striatum and motor cortex respectively. The pairwise spike correlations were remarkably stable, remaining essentially unchanged even after 10 days (Figure 4B-C), consistent with a very stable underlying network (Grutzendler et al., 2002; Yang et al., 2009).

**Figure 4:**
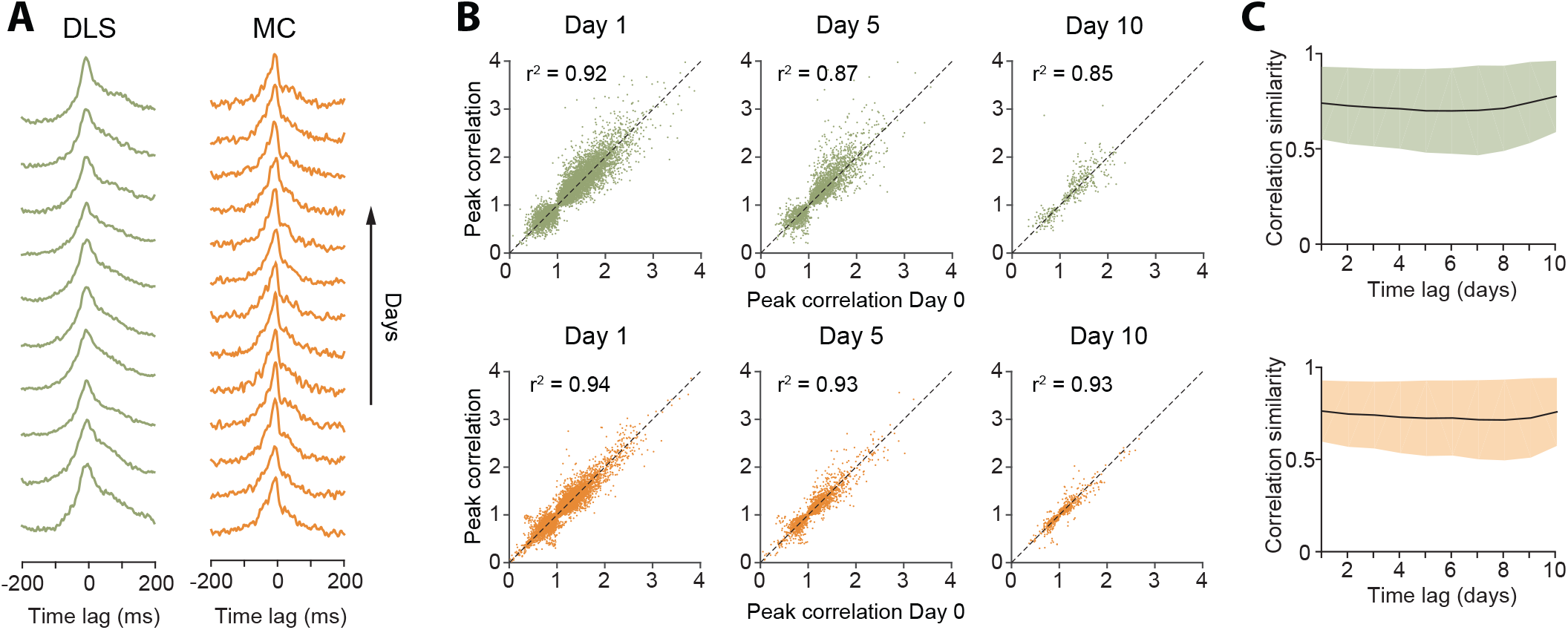
Stability of network dynamics A. Correlograms for example unit pairs from DLS (right) and MC (left) over 10+ days. Each line represents the normalized correlogram measured on a particular day.
B. Scatter plots comparing the peak correlation of a unit pair on a baseline day to the same pair’s peak correlation 1, 5 or 10 days later, for DLS (top) and MC (bottom). The black line indicates equality. Values >1 (or <1) correspond to positive (or negative) correlations (Supplemental Experimental Procedures). Day 1: n=10506 comparisons for DLS and n=17918 for MC. Day 5: n=3004 comparisons for DLS and n=4208 for MC. Day 10: n=390 comparisons for DLS and n=999 for MC. All unit pairs whose correlograms had at least 5000 spikes over a day’s period were included.
C. Stability of correlations over time. The correlation similarity (see Supplemental Experimental Procedures) was measured across time-lags of 1 to 10 days for pairs of units (solid lines) in DLS (top) and MC (bottom). Colored shaded regions indicate the standard deviation of the correlation similarity between all pairs that had significant correlograms on at least one recording day (Supplemental Experimental Procedures).

### Stability of behavioral state-dependent activity

Our results demonstrated long-term stability in the time-averaged statistics of single units (Figure 3) as well as their interactions (Figure 4), both in motor cortex and striatum. However, the brain functions to control behavior, and the activity of both cortical and striatal units can be expected to differ for different behaviors. To better understand the relationship between the firing properties of single units and ongoing behavior, and to assess the stability of these relationships, we analyzed single unit activity in different behavioral states. To do this, we developed an algorithm for automatically classifying a rat’s behavior into ‘states’ using highresolution measurements of head acceleration and local field potentials (LFPs). Our method distinguishes grooming, eating, active exploration, task engagement as well as quiet wakefulness, rapid eye movement and slow wave sleep (Gervasoni et al., 2004; Venkatraman et al., 2010) (Figure S5, Supplemental Experimental Procedures), and assigns ∼85% of the recording time to one of these states.

Behavioral state transitions, such as between sleep and wakefulness, have been shown to affect average firing rates (Evarts, 1964; Lee and Dan, 2012; Peña et al., 1999; Santhanam et al., 2007; Steriade et al., 1974; Vyazovskiy et al., 2009), inter-spike interval distributions (Mahon et al., 2006; Vyazovskiy et al., 2009), and neural synchrony (Gervasoni et al., 2004; Ribeiro et al., 2004; Wilson and McNaughton, 1994). However, most prior studies analyzed brief recording sessions in which only a narrow range of behavioral states could be sampled (Mahon et al., 2006; Vyazovskiy et al., 2009; Wilson and McNaughton, 1994), leaving open the question of whether the modulation of neural dynamics across behavioral states is distinct for different neurons in a network and whether state-specific activity patterns of single neurons are stable over time. Not surprisingly, we found that firing rates of both striatal and motor cortical units depended on what the animal was doing (Figure 5A). Interestingly, the relative firing rates in different behavioral states (i.e. a cell’s ‘state tuning’) remained stable over weeks-long time scales (Figure 5A-B).

**Figure 5:**
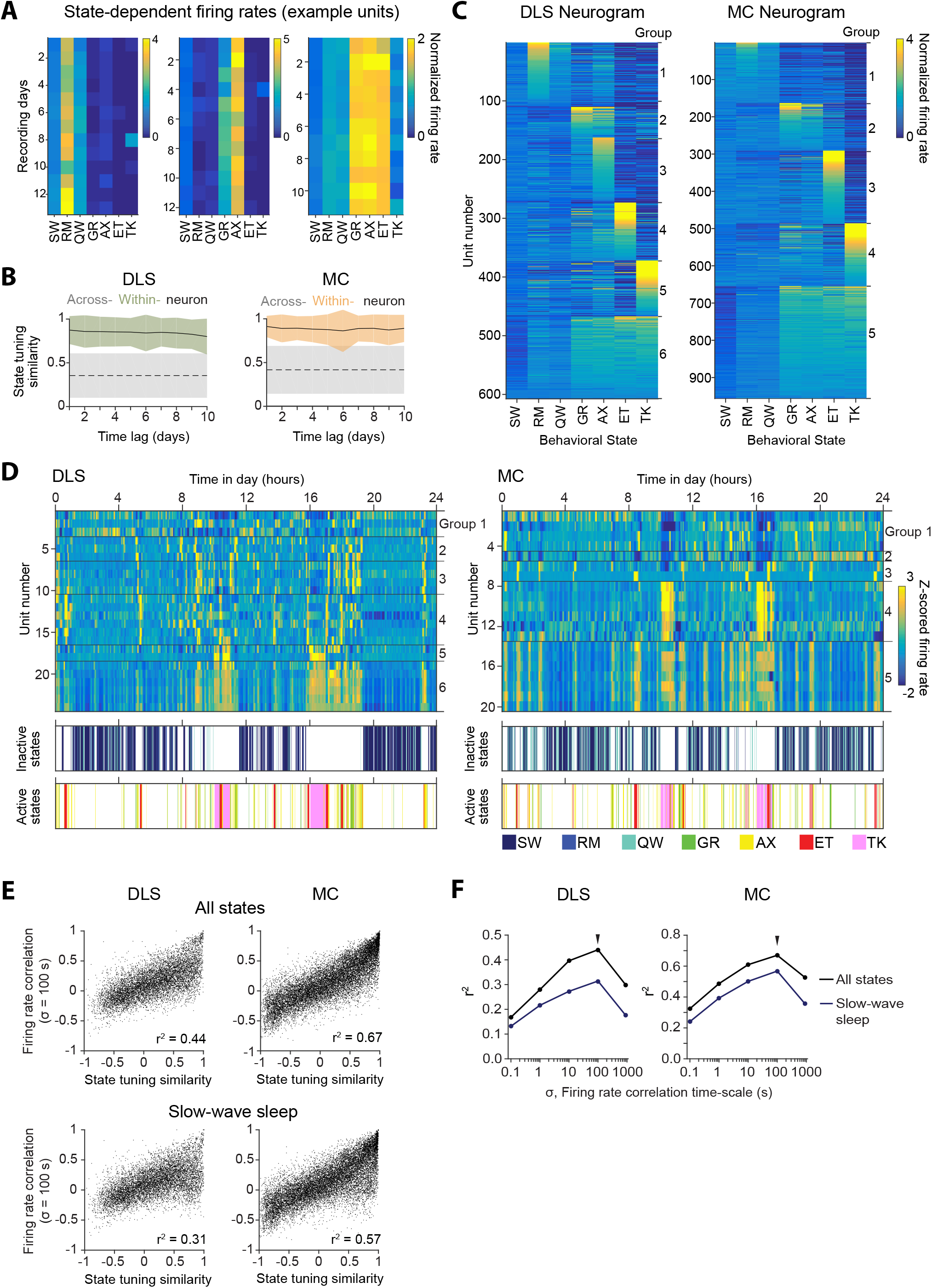
Stability of behavioral state-dependent activity patterns A. Stability of average firing rates in different behavioral states (i.e. the unit’s ‘statetuning’) across several days for example units recorded in the DLS (left) and MC (middle and right). The color-scale corresponds to the firing rate of a unit normalized by its average firing rate. The state abbreviations correspond to slow-wave sleep (SW), REM sleep (RM), quiet wakefulness (QW), grooming (GR), active exploration (AX), eating (ET) and task execution (TK).
B. Stability of units’ state-tuning over time. Correlations between the state-tuning profiles (as in ‘A’) were measured across time-lags of 1 to 10 days for the same unit (within-unit, solid lines), or between simultaneously recorded units (across-unit, dashed lines) in MC (left) and DLS (right). Colored shaded regions indicate the standard deviation of within-unit state-tuning similarity, over all units. Grey shaded regions indicate standard deviation of across-unit state-tuning similarity, over all time-bins.
C. Clustering of units into functional ‘groups’ based on similarities in their state-tuning for units in DLS (left, 6 groups) and MC (right, 5 groups). Each row represents the average firing rate modulation across behavioral states for a particular unit, normalized by its average firing rate. Units are clustered into groups by k-means clustering, and sorted, within types, by their maximum normalized firing rates.
D. Circadian profiles of instantaneous firing rates (top) for simultaneously recorded DLS (left) and MC (right) units. Each row represents the Z-scored firing rate of a unit over the same 24 hour period. Units are vertically sorted by membership in functional groups per panel C. Behavioral states are indicated below by the labeled color scheme.
E. Scatter plots of pairwise instantaneous firing rate correlations versus state-tuning similarity for DLS (left) and MC (right) units. Dots represent all simultaneously recorded pairs of units (n = 8166 DLS and 13905 MC unit pairs). Correlations were calculated for time-varying firing rates in all labeled states (top) or during slow-wave sleep alone (bottom). All instantaneous firing rates were smoothed with a Gaussian filter having a standard deviation (σ) of 100 ms.
F. Coefficient of determination (or variance explained, r^2^) for the regression analysis in panel ‘E’, as a function of timescale of instantaneous firing rate smoothing (σ) for all simultaneously recorded DLS (left) and MC (right) units. Firing rate correlations were computed over all labeled states (black line) or during slow-wave sleep alone (blue line). Arrows indicate timescale at which maximum correlation was observed (σ = 100 s), also shown in panel E.

Though our results revealed a rich diversity in the state tuning of neurons in striatum and motor cortex (Figure 5A), we found that they clustered into relatively few groups based on similarities in their state tuning (Figure 5C, Methods). Although the clustering was carried out independently for cortical and striatal populations, the ‘functional groups’ that emerged had similar state tuning profiles across the two brain areas (Figure 5C).

Interneuronal spike correlations were also a function of behavioral state (Figure S6A-B), and this dependence was also stable over time (Figure S6C). We next probed whether the functional groups we had identified based on similarity in state-tuning (Figure 5C) corresponded to Hebbian ‘cell assemblies’, i.e. neurons with high interneuronal spike correlations (Buzsáki, 2010; Harris, 2005; Hebb, 1949; Singer et al., 1997). Interestingly, functional group identity did not predict the strength of pairwise spike correlations (Figure S6D), suggesting that the groups of neurons that share state tuning profiles are distinct from groups of neurons (i.e. ‘cell assemblies’) with strong interneuronal correlations (Buzsáki, 2010; Harris, 2005).

Co-fluctuations in the firing rate of neurons (Bair et al., 2001; Freeman et al., 2014; Kato et al., 2015) or between compound neural activity in different brain regions (Biswal et al., 1995; Buckner et al., 2013) have been shown to occur over longer time scales (<0.1 Hz) also in other systems, including humans. Such slow fluctuations have been attributed to changes in subjects’ internal state (Bair et al., 2001; Buckner et al., 2013). Consistent with this, we found that neurons belonging to the same functional group had strikingly similar firing rate modulation both over time and across behavioral states (Figure 5D). To quantify this interaction, we regressed firing rate correlations against the similarity in behavioral state-tuning for pairs of units (Figure 5E, Supplemental Experimental Procedures). We found that firing rate correlations accounted for a significant fraction of the variance in state-tuning similarity (Figure 5E-F), particularly when the firing rate was averaged on a scale that matched the dominant state transition timescales (91.44 s for the DLS rat and 94.09 s for the MC rat, Supplemental Experimental Procedures). Moreover, firing rate correlations were far greater for unit pairs belonging to the same functional group than for pairs from different groups (Figure S6E).

To test whether co-fluctuations in the firing rate of neurons within a functional group (Figure 5D) are solely driven by behavioral state transitions, we repeated the above analysis (Figure 5E-F) within a single behavioral state – slow-wave sleep (the state where we had most data). Interestingly, the relationship between pairwise firing rate correlations in slow-wave sleep and pairwise state-tuning similarity remained strong (Figure 5E-F), suggesting that the functional groups represent more than simply groups of neurons tuned to the same behavioral state.

### Stability in the coupling between neural and behavioral dynamics

Neurons in motor-related areas encode and generate motor output, yet the degree to which the relationship between single unit activity and various movement parameters (i.e. a cell’s ‘motor tuning’) is stable over days and weeks has been the subject of recent debate (Carmena et al., 2005; Chestek et al., 2007; Ganguly and Carmena, 2009; Rokni et al., 2007; Stevenson et al., 2011). Long-term continuous recordings during natural and learned behaviors constitute unique datasets for characterizing the stability of movement coding at the level of single neurons.

We addressed this by computing the ‘response fields’ (Cheney and Fetz, 1984; Cullen et al., 1993; Serruya et al., 2002; Wessberg et al., 2000), i.e. the spike-triggered average (STA) accelerometer power (Supplemental Experimental Procedures), of neurons in different active states (Figure 6; Supplemental Experimental Procedures). The percentage of striatal units with significant response fields during grooming, exploration and eating was 17.1% (n=118), 16.0% (n=111) and 17.5% (n=121), respectively. In motor cortex, the numbers were higher: 28.8% (n=291), 23.6% (n=239) and 32.7% (n=331) for the three active states respectively. Movement STAs varied substantially between neurons and across behavioral states, but were stable for the same unit when compared over days within a particular state (Figure 6A-B).

**Figure 6:**
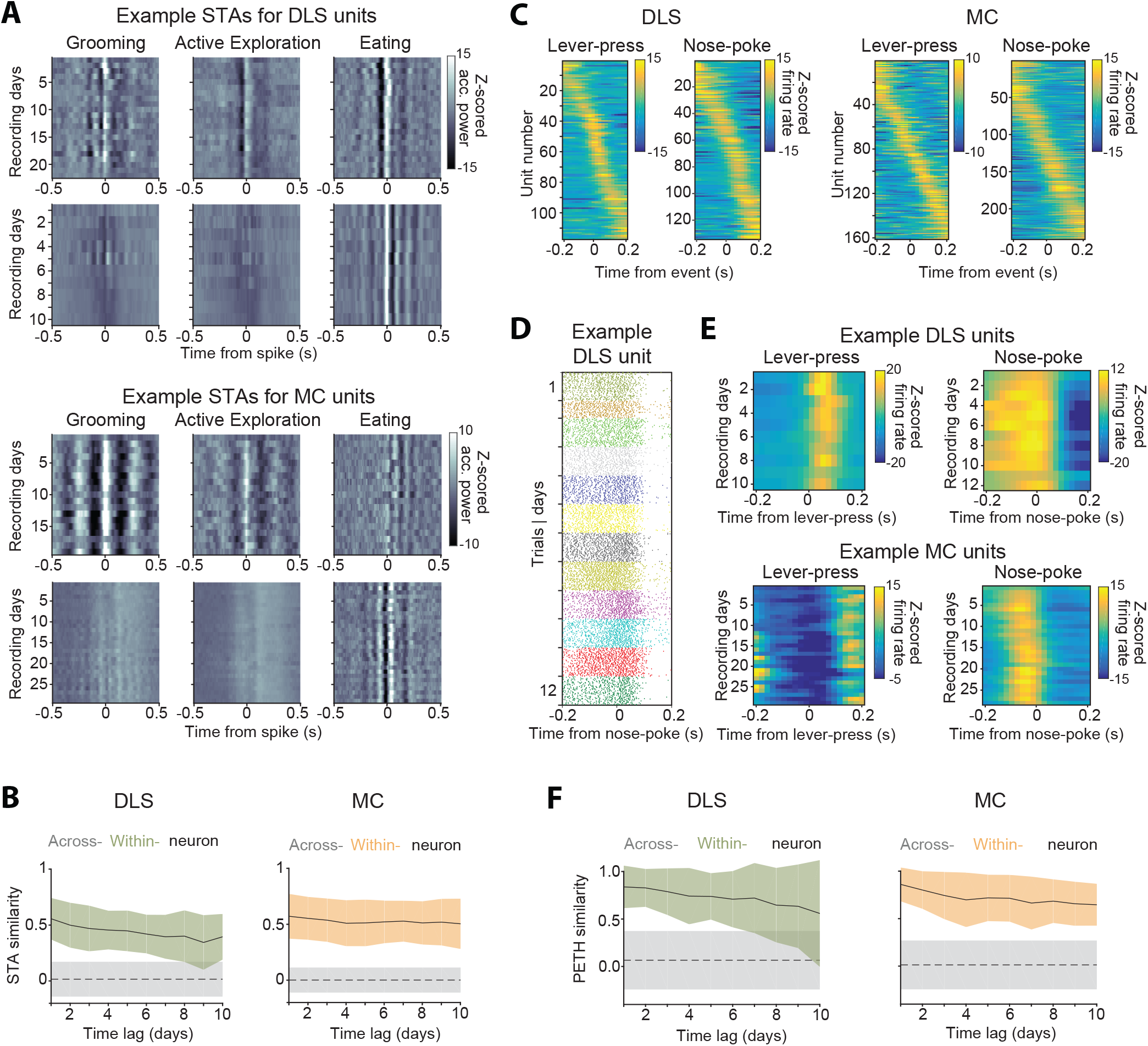
Stability of behavioral representations. A. Spike-triggered average (STA) accelerometer power calculated daily in three different behavioral states – grooming (left), active exploration (middle) and eating (right). Shown are four example units recorded from DLS (top two rows) and MC (bottom two rows).
B. Stability of STAs over time. Correlations between STAs were measured across time-lags of 1 to 10 days for the same unit (within-unit, solid lines), or between simultaneously recorded units (across-unit, dashed lines) in MC (left) and DLS (right). Colored shaded regions indicate the standard deviation of within-unit STA similarity (averaged over the 3 behavioral states in ‘A’), over all units. Grey shaded regions indicate standard deviation of across-unit STA similarity, over all time-bins.
C. Peri-event time histograms (PETHs) of DLS (left) and MC (right) unit activity, aligned to the timing of a lever-press or nose-poke during the execution of a skilled motor task. Plotted are the PETHs of units that had significant modulations in their firing rate in a time-window ±200 ms around the time of the behavioral event (Supplemental Experimental Procedures). The color scale indicates Z-scored firing rate. Units are sorted based on the times of peaks in their PETHs.
D. Spike raster of an example DLS unit over 12 days, aligned to the time of a nose-poke event. Each dot represents a spike-time on a particular trial. The color of the dot indicates day of recording.
E. PETHs computed over several days for example DLS (top) and MC (bottom) units to lever-press (left) and nose-poke (right) events in our task.
F. Stability of task PETHs over time. Correlations between PETHs were measured across time-lags of 1 to 10 days for the same unit (within-unit, solid lines), or between simultaneously recorded units (across-unit, dashed lines) in DLS (left) and MC (right). Colored shaded regions indicate the standard deviation of within-unit PETH similarity (averages across lever-press and nose-poke events), over all units. Grey shaded regions indicate standard deviation of across-unit PETH similarity (averaged across lever-press and nose-poke events), over all time-bins.

Another measure used to characterize neural coding of behavior on fast timescales is the ‘peri-event time histogram’ (PETH), i.e. the average neural activity around the time of a salient behavioral event. We calculated PETHs for two events reliably expressed during the execution of a skill our subjects performed in daily training sessions (Kawai et al., 2015) (Experimental Procedures): (i) the first lever-press in a learned lever-press sequence and (ii) entry into the reward port after successful trials (‘nose-poke’). The fraction of units whose activity was significantly locked to the lever-press or nose-poke (see Supplemental Experimental Procedures) was 16.9% (n=117) and 19.3% (n=134) in striatum, and 15.9% (n=161) and 23.8% (n=241) in motor cortex (Figure 6C), respectively. When comparing PETHs of different units for the same behavioral events, we observed that the time of peak firing was distributed across a range of delays relative to the time of the events (Figure 6C). However, despite the heterogeneity in PETHs across the population of units, the PETHs of individual units were remarkably similar when compared across days (Figure 6D-F).

To probe whether the functional groups we identified based on their state-tuning profiles (Figure 5C) reflect similar motor tuning, we assessed the similarity of movement STAs and PETHs of units between and across functional groups. We found no consistent relationship between motor tuning and behavioral state tuning (Figure S7), suggesting that functional groups reflect a higher order behavioral state-dependence, and not merely similar movement tuning.

## Discussion

Recording from populations of single neurons in behaving animals has traditionally been a laborious undertaking both in terms of experimentation and analysis. Automating this process, and extending the recordings over weeks and months, increases the efficiency and power of such experiments in multiple ways. First, it eliminates the manual steps of intermittent recordings, such as transferring animals to and from their recording chambers, plugging and unplugging recording cables etc., which, besides being time-consuming, can be detrimental to the recordings (Buzsáki et al., 2015) and stressful for the animals (Dobrakovová and Jurcovicová, 1984; Longordo et al., 2011). Second, when combined with fully automated home-cage training (Poddar et al., 2013), this approach allows the neural correlates of weeks- and months-long learning processes to be studied in an efficient manner. Indeed, our system should enable a single researcher to supervise tens, if not hundreds, of recordings simultaneously, thus radically improving the ease and throughput with which such experiments can be performed. Third, continuous recordings allow the activity of the same neurons to be tracked over days and weeks, allowing their activity patterns to be pooled and compared for more trials, and across different experimental conditions and behavioral states, thus increasing the power with which the principles of neural function can be inferred (Lütcke et al., 2013).

Despite their many advantages, continuous long-term neural recordings are rarely performed. This is in large part because state-of-the-art methods for processing extracellular recordings require significant human input and hence make the analysis of large datasets prohibitively time-consuming. Our fully automated spike tracking (FAST) algorithm overcomes this bottleneck, thereby dramatically increasing the feasibility of continuous and high-throughput long-term recordings of extracellular signals in behaving animals.

### Long-term stability of neural dynamics

Recording the activity of the same neuronal population over long time periods can be done by means of functional calcium imaging (Huber et al., 2012; Lütcke et al., 2013; Peters et al., 2014). However, the calcium signal reports a relatively slow correlate of neural activity that can make it difficult to reliably resolve individual spike events and hence to assess fine timescale neuronal interactions (Grienberger and Konnerth, 2012; Lütcke et al., 2013). Furthermore, calcium indicators function as chelators of free calcium ions (Hires et al., 2008; Tsien, 1980) and could interfere with calcium-dependent plasticity processes (Zucker, 1999) and, over the long-term, compromise the health of cells (Looger and Griesbeck, 2012). These variables may have contributed to discrepancies in studies using longitudinal calcium imaging, with some reporting dramatic day-to-day fluctuations in neural activity patterns (Huber et al., 2012) while others report more stable representations (Peters et al., 2014), leaving open the question of how stable the brain really is.

While electrical measurements of spiking activity do not suffer from the same drawbacks, intermittent recordings may fail to reliably track the same neurons over multiple days (Figure S4D-E) (Dickey et al., 2009; Emondi et al., 2004; Fraser and Schwartz, 2012; Tolias et al., 2007). Attempts at inferring long-term stability of movement-related neural dynamics from such experiments have produced conflicting results, with some studies reporting stability (Chestek et al., 2007; Fraser and Schwartz, 2012; Ganguly and Carmena, 2009; Greenberg and Wilson, 2004), while others finding significant fluctuations even within a single day (Carmena et al., 2005; Rokni et al., 2007). Such contrasting findings could be due to systematic errors in unit tracking across discontinuous recording sessions (Figure S4D-E).

Our continuous recordings, which allow for the reliable characterization of single neuron properties over long time–periods, suggest a remarkable stability in both spiking statistics (Figure 3), neuronal interactions (Figure 4), and tuning properties (Figures 5,6) of motor cortical and striatal units. Taken together with studies showing long-term structural stability at the level of synapses (Grutzendler et al., 2002; Yang et al., 2009), our findings suggest that neural networks are, over all, highly stable systems, both in terms of their dynamics and connectivity.

### Neuronal ensembles defined by behavioral states

An intriguing finding emerging from our analysis of continuous long-term recordings was that neurons clustered into a few distinct groups based on similarities in firing rate modulation across behavioral states (Figure 5C). Group membership is not simply a consequence of units having similar motor tuning (Figure S7) (Ganguly and Carmena, 2009), nor is it consistent with synchronization of spiking activity, as in previously described ‘cell assemblies’ (Buzsáki, 2010; Harris, 2005; Hebb, 1949). Instead, the functional groups reflect co-fluctuations in the firing rate of neurons on timescales corresponding to average transition times between behavioral states (∼100 s). Such functional groups have been described in other systems, including networks of neurons identified by whole-brain calcium imaging studies in nematodes (Kato et al., 2015) and zebrafish (Ahrens et al., 2013; Freeman et al., 2014), as well as networks of functionally connected areas in the human brain identified by fMRI (Biswal et al., 1995; Buckner et al., 2013).

Slow, correlated fluctuations in the activity of forebrain circuits (Leopold et al., 2003) are thought to be driven by ascending neuromodulatory input from brainstem and hypothalamic nuclei involved in behavioral state transitions (Lee and Dan, 2012). If these projections are responsible for the firing rate correlations we observe, it would imply that neurons in different functional groups are differentially modulated by them. Different cell types can respond differently to behavioral state transitions (Gentet et al., 2010), suggesting that functional groups may reflect distinct cell-types. Future experiments involving continuous long-term recordings in combination with optogenetic cell-type identification (Lima et al., 2009) and perturbations of neuromodulatory pathways (Adamantidis et al., 2007; Carter et al., 2010) will be able to inform this question.

## Future improvements

There is a continued push to improve recording technology to increase the number of neurons that can be simultaneously recorded from (Berényi et al., 2014; Buzsáki et al., 2015; Du et al., 2011; Viventi et al., 2011), as well as make the recordings more stable (Alivisatos et al., 2013; Chestek et al., 2011; Cogan, 2008; Felix et al., 2013; Guitchounts et al., 2013; Zhou et al., 2014). Our modular and flexible experimental platform can easily incorporate such innovations. Importantly, our novel and automated analysis pipeline (FAST) solves a major bottleneck downstream of these solutions by allowing increasingly large datasets to be efficiently parsed, thus making automated recording and analysis of large populations of neurons in behaving animals a feasible prospect.

## Experimental Procedures

### Animals

The care and experimental manipulation of all animals were reviewed and approved by the Harvard Institutional Animal Care and Use Committee. Experimental subjects were female Long Evans rats, 3-8 months old at the start of the experiment (n=2, Charles River).

### Behavioral training

Before implantation, rats were trained twice daily on a timed lever-pressing task (Kawai et al., 2015) using our fully automated home-cage training system (Poddar et al., 2013). Once animals reached asymptotic performance on the task, they underwent surgery to implant tetrode drives into dorsolateral striatum (n=1) and motor cortex (n=1) respectively (see Supplementary Experimental Procedures). After 7 days of recovery, rats were returned to their home-cages, which had been outfitted with an electrophysiology recording extension (Figure 1A, Supplementary Experimental Procedures). The cage was placed in an acoustic isolation box, and training on the task resumed. Neural and behavioral data was recorded continuously for 12-16 weeks with only brief interruptions (median time of 0.2 hours) for occasional troubleshooting.

### Continuous behavioral monitoring

To monitor the rats’ head movements continuously, we placed a small 3-axis accelerometer (ADXL 335, Analog Devices) on the recording head-stage. The output of the accelerometer was sampled at 7.5 kHz per axis. We also recorded 24/7 continuous video at 30 frames per second with a CCD camera (Flea 3, Point Grey) or a webcam (Agama V-1325R). Video was synchronized to electrophysiological signals by recording TTL pulses from the CCD cameras that signaled frame capture times.

### Surgery

Rats underwent surgery to implant a custom-built recording device (see ‘Tetrode arrays’). Animals were anesthetized with 1-3% isoflurane and placed in a stereotax. The skin was removed to expose the skull and five bone screws (MD-1310, BASi), including one soldered to a 200 μm diameter silver ground wire (786500, A-M Systems), were driven into the skull to anchor the implant. A stainless-steel reference wire (50 μm diameter, 790700, AM-Systems) was implanted in the external capsule to a depth of 2.5 mm, at a location posterior and contralateral to the implant site of the electrode array. A 4-5 mm diameter craniotomy was made at a location 2 mm anterior and 3 mm lateral to bregma for targeting electrodes to motor cortex, and 0.5 mm anterior and 4 mm lateral to bregma for targeting dorsolateral striatum. After removing the dura-mater, the pia-mater surrounding the implant site was glued to the skull with cyanoacrylate glue (Krazy glue). The pia-mater was then weakened using a solution of 20 mg/ml collagenase (Sigma) and 0.36 mM calcium chloride in 50 mM HEPES buffer (pH 7.4) in order to minimize dimpling of the brain surface during electrode penetration (Kralik et al., 2001). The 16-tetrode array was then slowly lowered to the desired depth of 1.85 mm for motor cortex and 4.5 mm for striatum. The microdrive was encased in a protective shield and cemented to the skull by applying a base layer of Metabond luting cement (Parkell) followed by dental acrylic (A-M systems).

### Histology

At the end of the experiments, animals were anesthetized and anodal current (30 μA for 30 s) passed through select electrodes to create micro-lesions at the electrode tips. Animals were transcardially perfused with phosphate-buffered saline (PBS) and subsequently fixed with 4% paraformaldehyde (PFA, Electron Microscopy Sciences) in PBS. Brains were removed and postfixed in 4% PFA. Coronal sections (60 μm) were cut on a Vibratome (Leica), mounted, and stained with cresyl violet to reconstruct the location of implanted electrodes. All electrode tracks were consistent with the recordings having been done in the targeted areas.

### Tetrode arrays

Tetrodes were fabricated by twisting together short lengths of four 12.5 μm diameter nichrome wires (Redi Ohm 800, Sandvik-Kanthal), after which they were bound together by melting their polyimide insulation with a heat-gun. An array of 16 such tetrodes was attached to a custom-built microdrive, advanced by a 0-80 threaded screw (∼320 μm per full turn). The wires were electroplated in a gold (5355, SIFCO) and 0.1% carbon nanotubes (Cheap Tubes dispersed MWNTs, 95wt% <8 nm) solution with 0.1% polyvinylpyrrolidone surfactant (PVP-40, Sigma-Aldrich) to lower electrode impedances to ∼100-150 kΩ (Ferguson et al., 2009; Keefer et al., 2008). The prepared electrode array was then implanted into the motor cortex or striatum.

### Automatic classification of behavior

We developed an unsupervised algorithm to classify behaviors based on accelerometer data, LFPs, spike times, and task event times (Gervasoni et al., 2004; Venkatraman et al., 2010). Behaviors were classified at 1 s resolution into one of the following ‘states’: grooming (GR), eating (ET), active exploration (AX), task engagement (TK), quiet wakefulness (QW), rapid eye movement sleep (RM) or slow-wave sleep (SW). See Supplementary Experimental Procedures for detailed algorithm.

### Analysis of Neural Data

*Automated spike sorting*. We developed a new and automated spike sorting algorithm that is able to efficiently track single units over long periods of time. Our algorithm first identifies the spikes from the raw recordings and then performs two fully automated processing steps. See Supplemental Experimental Procedures for the detailed algorithm.

All subsequent analysis of neural data was carried out using custom scripts in Matlab (Mathworks). See Supplemental Experimental Procedures for detailed description of the analyses.

## Acknowledgements.

This work was supported by a McKnight Scholars Award (BPÖ), HSFP and EMBO fellowships (SBEW), and by the Swartz Foundation (EK). AKD is an Ellison Medical Foundation fellow of the Life Sciences Research Foundation. We thank Michelle Choi for help with behavioral scoring from videos of rats.

## Author Contributions

BPÖ, RP and AKD designed the study with input from all authors. RP designed and implemented the algorithm for automated spike sorting with the help of AKD and VN. AKD performed the experiments. AKD analyzed the neural data with help from BPÖ and SBEW. EK wrote the software for automated behavioral analysis and analyzed the behavior together with AKD. BPÖ, RP and AKD wrote the manuscript.

---

**Figure S1:**
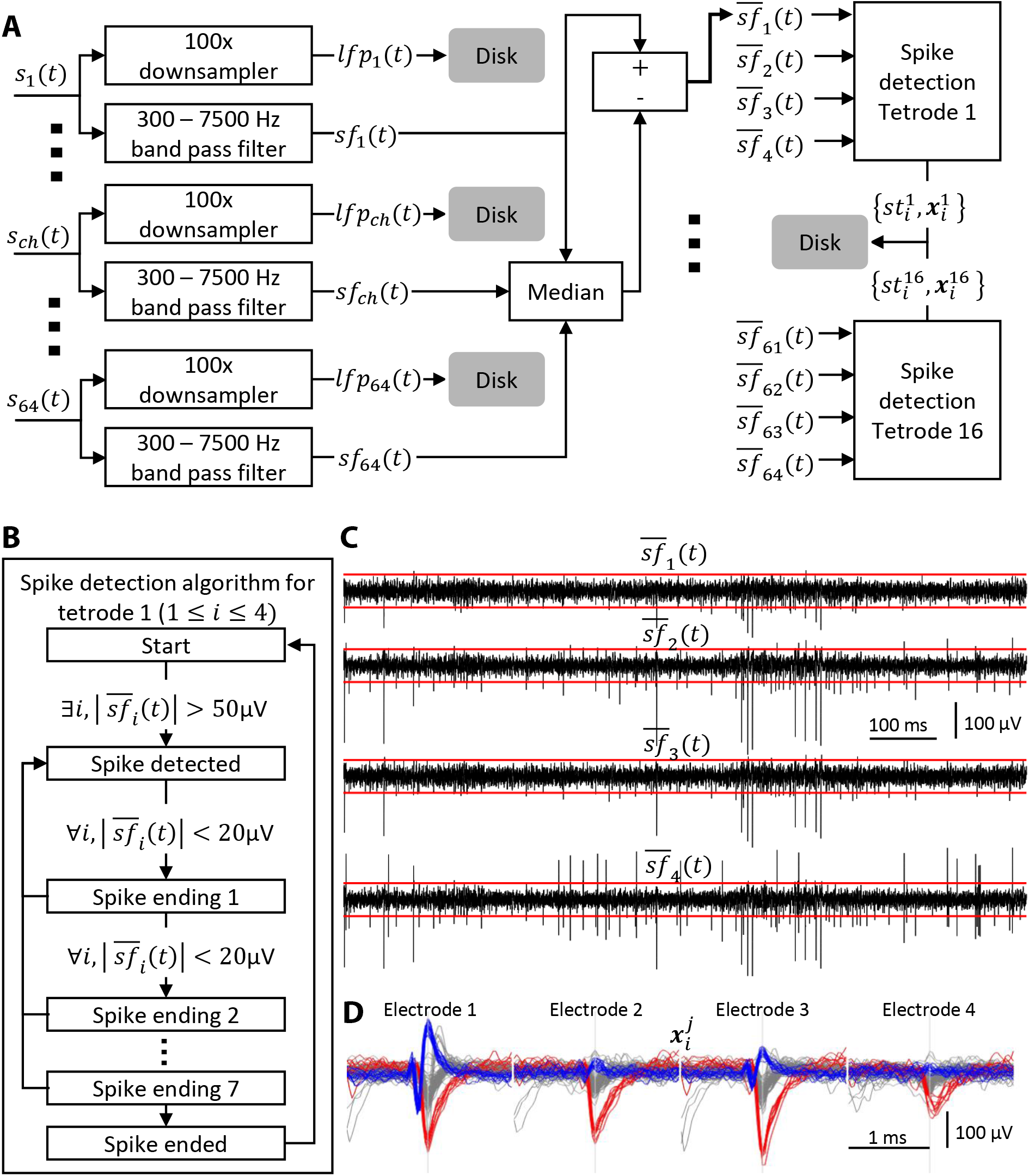
Algorithm for identifying spikes from tetrode recordings. A. Each input channel *s_ch_* is split into two streams, one containing the low frequency component *lfp_ch_* and one containing the high frequency one, *sf_ch_*. The median of *sf_ch_* across all channels is subtracted from each channel resulting in 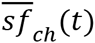. Spike times 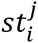 and spike waveforms 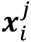 from each tetrode are then extracted. The LFPs and spikes extracted from the raw data are saved to disk resulting in a 5-10x ‘compression’ of the raw data.
B. Algorithm for detecting spikes. If the absolute value of the filtered signal exceeds 50 μV in any channel of a tetrode then a spike is ‘detected’. The spike is considered to have ‘ended’ if all channels remain within 20 μV for 8 consecutive samples.
C. Example of a 1 s long raw recording from a tetrode. The red lines mark the ±50 μV spike detection threshold.
D. Examples of 2.13 ms wide spike snippets (64 samples) extracted from the data in **C**. Snippets from all 4 electrodes detected using the state machine in **B** are aligned to the peak of the spike waveform and concatenated to produce the 256 sample spike waveforms 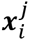.

**Figure S2:**
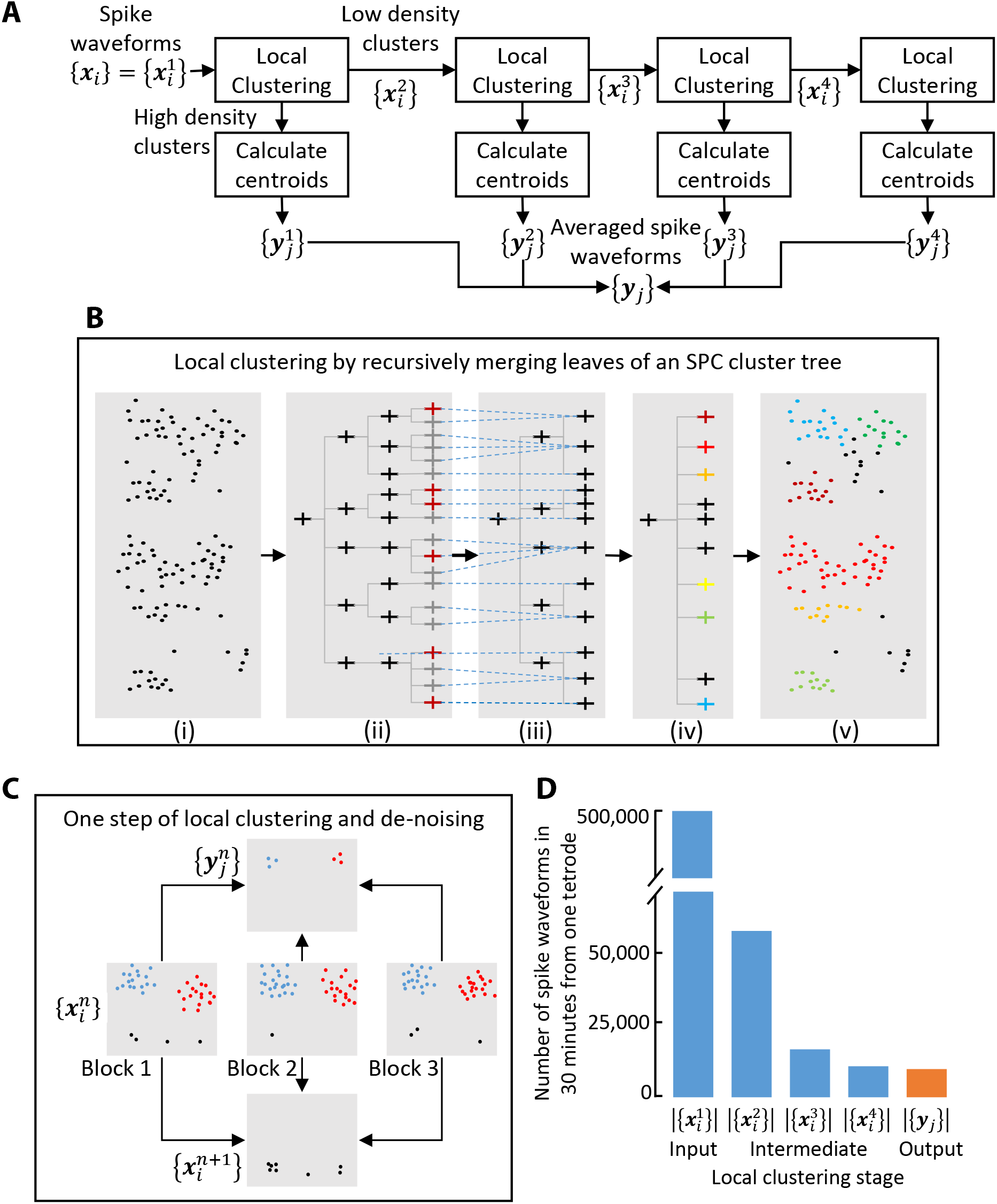
Algorithm for local clustering and de-noising. A. Raw spike waveforms {*x_i_*} are locally clustered and split into low- and high-density clusters (details in panels B and C). The spikes from low-density clusters are further split into two streams in the same manner 3 more times. The centroids of high density clusters from all 4 stages are pooled together to form the output {*y_j_*}.
B. Local clustering of each 1000 spike block. Super-paramagnetic clustering generates a cluster tree (ii) from the spike waveforms (i), the leaves of which are recursively merged (iii and iv) to generate a clustering of the 1000 points (v). The dotted blue lines show which leaves of the tree in (ii) are merged to produce the tree in (iii). The nodes marked red in (ii) correspond to ‘distinct clusters’, i.e. clusters that are very different from the parent nodes. The leaves of (iii) are similarly merged to produce the tree in (iv). The colored leaves correspond to high-density clusters, i.e. clusters with more than 15 points and the black leaves correspond to low-density clusters.
C. Schematic illustrating splitting of spikes into low-density and high-density clusters. The set of input spike waveforms 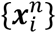 is split into blocks of 1000 spikes (3 blocks shown in the figure) with each block split into low (colored black) and high density clusters (colored blue and red) using the procedure shown in panel **B**. The spikes from the low density clusters are pooled together to form 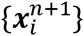. The centroids of the high density clusters form 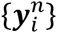.
D. Number of spike waveforms in a 30 minute period from one tetrode in various stages of the local clustering and de-noising algorithm.

**Figure S3:**
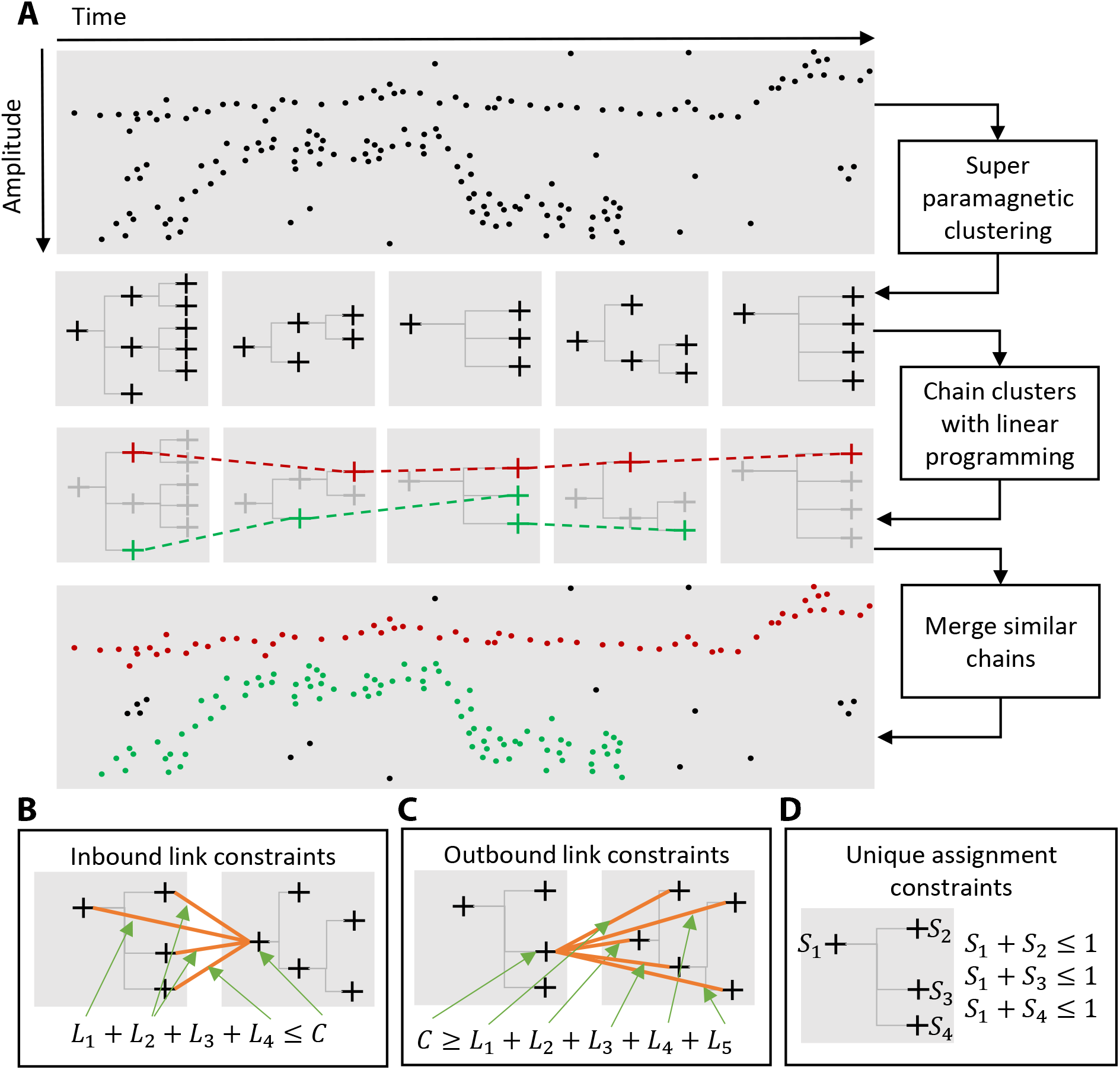
Algorithm for linking cluster trees to track single units over time. A. The output of the previous step (averaged spikes waveforms – see Figure S2) is split into a sequence of 1000 spike blocks and converted into a sequence of cluster trees (5 trees shown in the figure). A subset of all possible links between adjacent cluster trees is chosen by maximizing the total similarity between linked nodes subject to the constraints depicted in panels **B**, **C**, and **D**. The subset of chosen nodes and links are highlighted in color. Three sets of nodes connected by links, one in red and two in green, are shown. The two green chains are merged based on waveform similarity to produce a final sorting containing two units (red and green).
B. The constraint shown ensures that none of the 4 incoming links *[L_1_ – L_4_)* are chosen if the node marked *C* is not chosen. It also ensures that if *C* is chosen, at most one of the incoming links is chosen.
C. Same as **B** but for outgoing links.
D. These constraints ensure that if a node is chosen then none of its parents or child nodes are.

**Figure S4:**
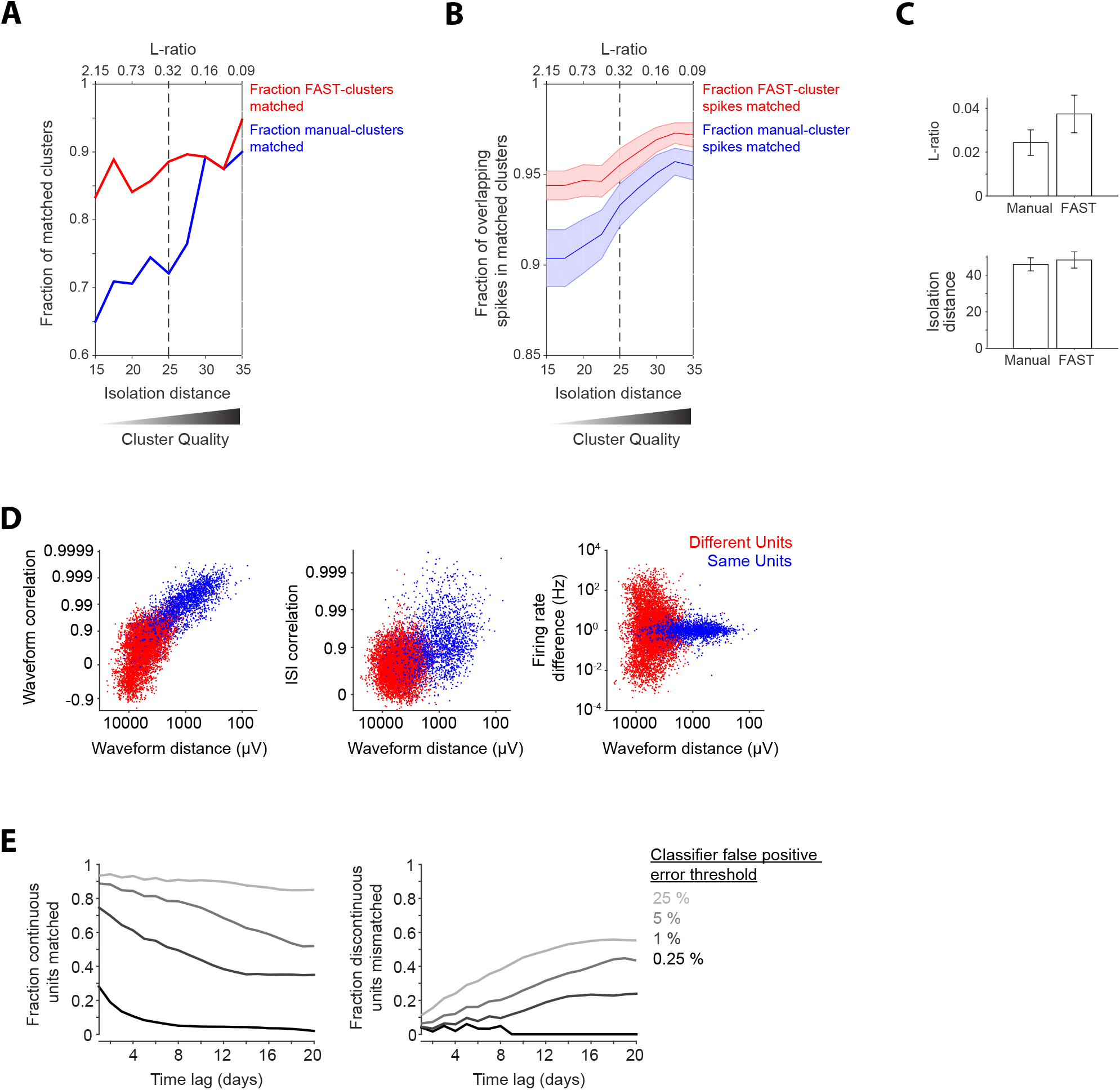
Benchmarking the performance of FAST by comparing its output to manual sorting and to discontinuous tracking of units across daily recording sessions. A. Fraction of clusters matched after independent spike sorting of the same dataset by manual sorting and our unsupervised sorting algorithm (FAST). The number of matched clusters (defined as sharing more than 50% of the same spikes) after sorting by the two methods is shown as a fraction of the total number of manual clusters (blue) or clusters identified by FAST (red). The x-axes indicate different cluster quality criteria, in terms of isolation distance (bottom) and L-ratio (top), used to eliminate low-quality clusters from both auto and manual sorting results. The dotted line indicates the quality criteria typically used in our recordings.
B. Fraction of spikes overlapping in matched clusters identified by both manual sorting and FAST. The number of overlapping spikes is shown as a fraction of total spikes in the manual cluster (blue) or the cluster identified by FAST (red). The x-axes indicate different cluster quality criteria, in terms of isolation distance (bottom) and L-ratio (top), used to eliminate low-quality clusters from the output of both FAST and manual sorting. The dotted line indicates the quality criteria typically used in our recordings. Shaded regions indicate standard error of the mean, across clusters.
C. Comparison of cluster quality, measured in terms of L-ratio (top) and isolation distance (bottom), for clusters matched between manual sorting and FAST, after eliminating low quality clusters using our typical quality criteria (dotted lines in A-B). Error bars indicate standard error of the mean.
D. Discontinuous tracking of units across daily recording sessions is error-prone. Similarity metrics between units isolated on consecutive days in hour-long recording sessions (data from the DLS rat). Three out of six possible combinations of similarity features are presented here – spike waveform correlation (left), inter-spike interval histogram (ISI) correlation (middle) and firing rate difference (right) versus spike waveform Euclidean distance. Each dot represents the similarity between pairs of units recorded on the same tetrode and classified as the ‘same’ (blue dot) or ‘different’ (red dot) by our FAST algorithm.
E. (Left) Percentage of ‘same’ unit pairs identified by the FAST method that were successfully matched by discontinuous tracking over time-lags ranging from 1 to 20 days. (Right) Percentage of discontinuously matched unit pairs that are classified as ‘different’ by FAST for time-lags ranging from 1 to 20 days. Graded colors correspond to different false positive rate thresholds of the ‘same’ versus ‘different’ linear classifier (see Supplemental Experimental Procedures) ranging from 0.25% (light gray) to 25% (black).

**Figure S5:**
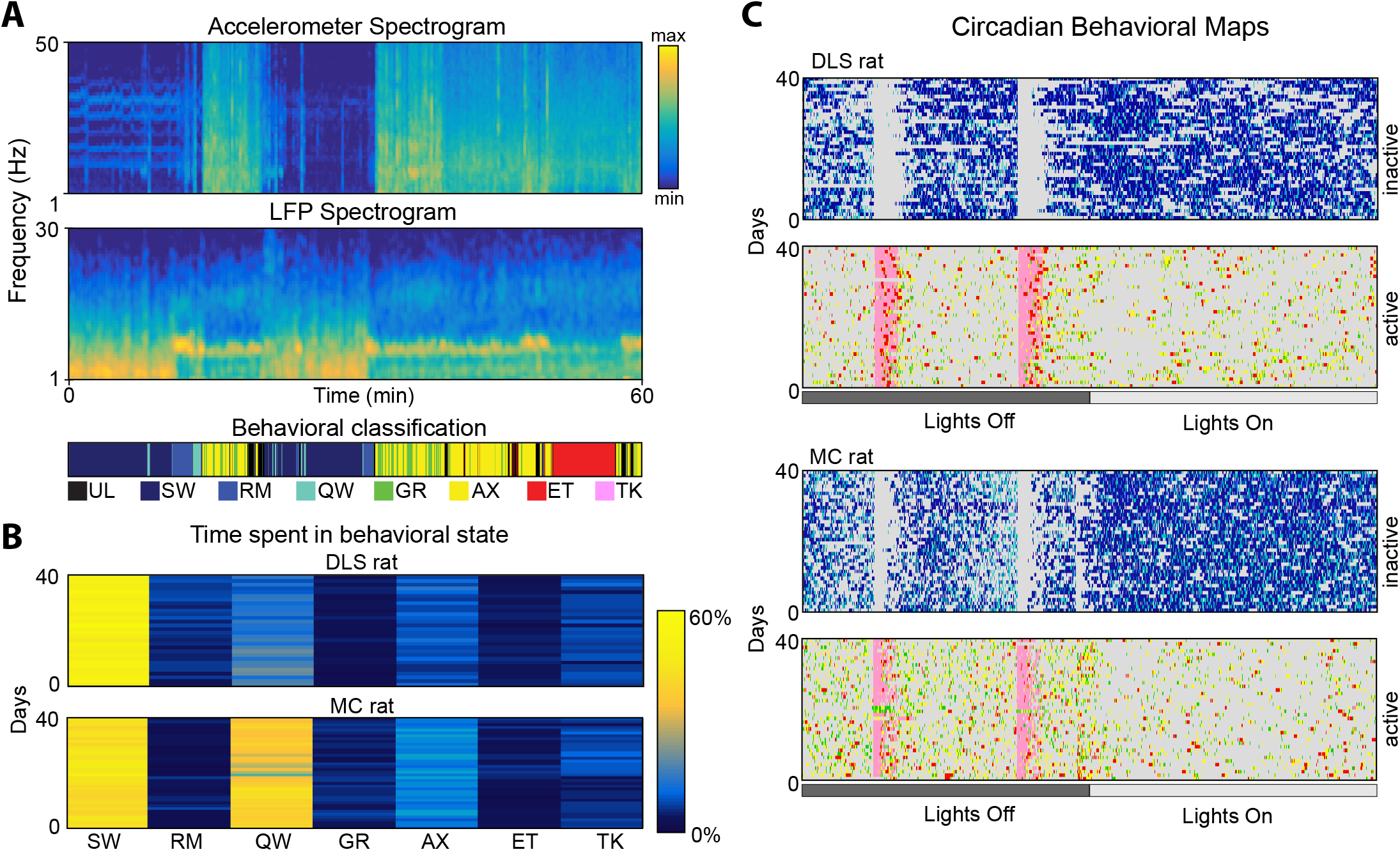
Automated classification of behavioral states. A. (Top) Spectrograms of the accelerometer signal (top) and the LFP (middle) for an hour-long window of recording from the striatum. Below is the color-coded output of our automated classification algorithm. UL-unlabeled; SW-Slow wave sleep; RM-REM sleep; QW-quiet wakefulness, GR-grooming, AX-active exploration, ET-eating, TK-task.
B. Ethograms showing the proportion of time spent in each of the behavioral states over 40 consecutive days for the two rats we recorded from. The ethograms do not include absence seizure-like states or unlabeled states and were normalized to one (Supplemental Experimental Procedures).
C. Circadian profiles of behavioral states over 40 days of recording for active and inactive states for the two rats in B. Behavioral states are color-coded as in A, with grey representing inactive (or active) and unlabeled states.

**Figure S6:**
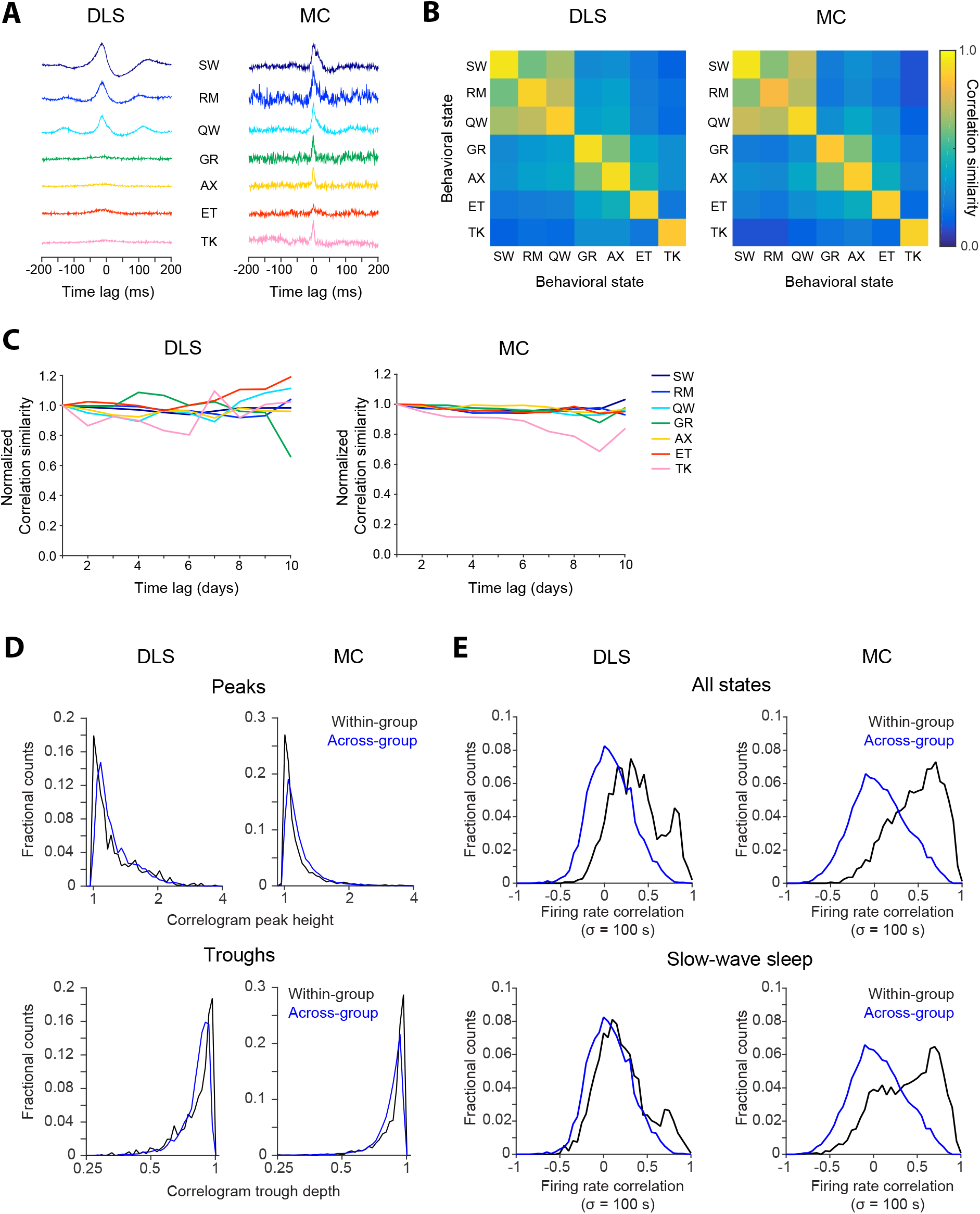
Network dynamics differ across behavioral states A. Cross-correlograms computed for example pairs of units in the DLS (left) and MC (right) in different behavioral states. Line colors represent behavioral state. The state abbreviations correspond to slow-wave sleep (SW), REM sleep (RM), quiet wakefulness (QW), grooming (GR), active exploration (AX), eating (ET) and task execution (TK).
B. Similarity of cross-correlograms measured in different behavioral states in DLS (left) and MC (right). The color-scale represents correlation similarity (see Supplemental Experimental Procedures). For each behavioral state, the ‘self’ similarity of correlations (corresponding to the diagonal of each plot) is computed after splitting the spikes recorded on alternate days into two correlograms and measuring the correlation coefficient between them.
C. Stability of cross-correlograms measured within specific behavioral states. The correlation similarity (see Supplemental Experimental Procedures) was measured across time-lags of 1 to 10 days (solid lines) in DLS (left) and MC (right) and normalized to its 1 day lag value. Line colors represent correlation similarity for different behavioral states, as per the plot legend.
D. Distributions of spike-train cross-correlogram peak heights (top) and trough depths (bottom) for simultaneously recorded DLS (left) and MC (right) unit pairs. Black lines correspond to pairs of units from the same functional group, while blue lines indicate units from different functional groups, as defined in Figure 5C. DLS n= 869 within-group and n=3520 across-group unit pairs; MC n=2962 within-group and n=8752 across-group unit pairs.
E. Distributions of pairwise firing rate correlations during all behavioral states (top) or slow-wave sleep (bottom) for all simultaneously recorded DLS (left) and MC (right) unit pairs. Black lines indicate pairwise firing rate correlations for units belonging to the same ‘functional group’ (see Figure 5C), while blue lines indicate units belonging to different functional groups. Firing rates were smoothed with a Gaussian having standard deviation σ = 100 s before computing the correlations. DLS n=1481 within-group and 6685 across-group unit pairs; MC n=3385 within-group and 10522 across-group unit pairs.

**Figure S7:**
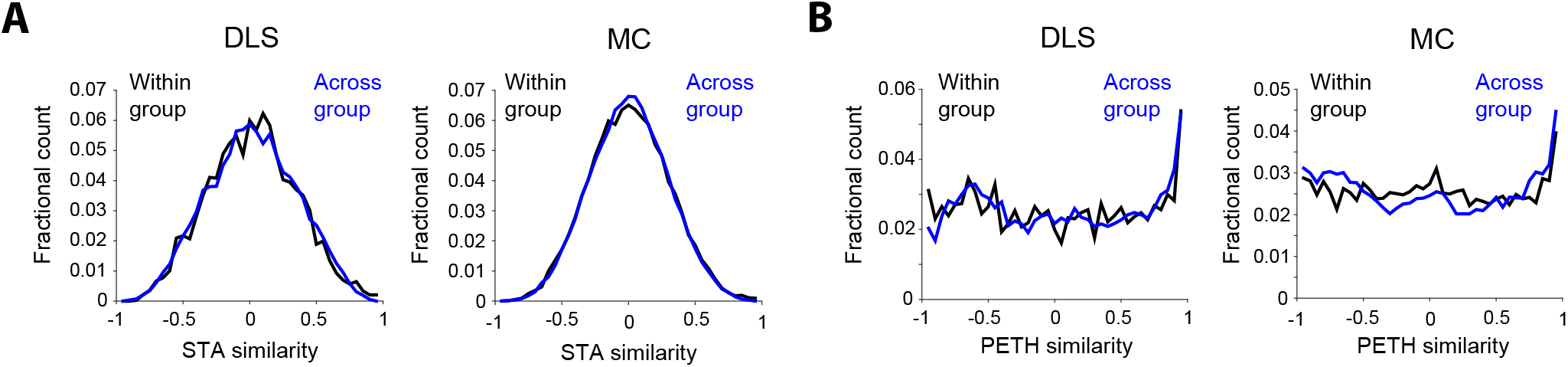
Similarity of motor representations in neurons with similar state-tuning. A. Histogram of STA similarity between all pairs of neurons belonging to the same (black) or different functional groups (blue) in the DLS (left, 6 groups) and MC (right, 5 groups). Clustering of neurons into groups is based on similarity in their state-tuning as shown in Figure 5C.
B. Histogram of PETH similarity between all pairs of neurons belonging to the same (black) or different functional group (blue) in the DLS (left, 6 groups) and MC (right, 5 groups).

## Supplemental Experimental Procedures

### Experimental infrastructure for long-term neural recordings in behaving animals

*Tethering system*. We used extruded aluminum with integrated V-shaped rails for the linear-slide (Inventables, Part #25142-03), with matching wheels (with V-shaped grooves) on the carriage (Inventables, Part #25203-02). The bearings in the wheels were degreased and coated with WD-40 to minimize friction. The carriage plate was custom designed (design files available upon request) and fabricated by laser cutting 3 mm acrylic. A low-cost 15-channel commutator (SRC-022, Hangzhou-Prosper Ltd) was mounted onto the carriage plate. The commutator was suspended with a counterweight using low friction pulleys (Misumi Part # MBFN23-2.1). We used a 14 conductor cable (Mouser, Part # 517-3600B/14100SF) to connect our custom designed head-stage to the commutator. The outer PVC insulation of the cable bundle was removed to reduce the weight of the cable and to make it more flexible. The same cable was used to connect the commutator to a custom designed data acquisition system. This cable needs to be as flexible as possible to minimize forces on the animals head.

*Recording hardware*. Our lightweight, low-noise and low-cost system for multi-channel extra-cellular electrophysiology uses a 64-channel head-stage (made of 2 RHD2132 ICs from Intantech), weighs less than 4 grams and measures 18 mm x 28 mm in size. The head-stage filters, amplifies, and digitizes extracellular neural signals at 16 bits/sample and 30 kHz per second per channel. An FPGA (Opal Kelly XEM6010) interfaces the head-stage with a computer that stores and displays the acquired data. Since the neural signals are digitized at the head-stage, the system is low-noise and can support long cable lengths. Design files and a complete parts list for all components of the system are available on request. The parts for a complete 64-channel electrophysiology extension, including head-stage, FPGA board, commutator, and pulley system, but excluding the recording computer, cost less than $1500.

### Analysis and automatic classification of behavior

*Processing of accelerometer data:* We performed wavelet analysis on the accelerometer signals to reconstruct a 5^th^ level approximation. This reconstruction, which captures information related to slow movements (similar to low pass filtering), was subtracted from the original signal. Accelerometer power was calculated in 1 ms bins from the 0-50 Hz frequency band. The power had a bimodal distribution with one narrow peak at low values corresponding to immobility and a secondary broad peak corresponding to epochs of movement. The threshold for immobility was set to best discriminate the two peaks.

*Processing of the LFP:* We computed the spectrogram for the LFP signal on each electrode in 1 s bins. The delta (1-4 Hz) and theta (5-9 Hz) power was normalized by the total power, averaged over all 64 channels, and smoothed with a 10 s boxcar filter. The delta power had a bimodal distribution with a broad peak around the low values corresponding to awake states and a secondary narrow peak with higher values corresponding to epochs of immobility. The threshold for high delta values was set to best discriminate the two peaks and was used to identify SWS epochs. Similarly, the theta power had a bimodal distribution with a narrow peak at low values corresponding to REM sleep states and a secondary broad peak at higher values (Eschenko et al., 2006; Gervasoni et al., 2004). The threshold for high theta values was set to best discriminate the two peaks and was used to identify REM epochs (Eschenko et al., 2006; Gervasoni et al., 2004).

*Classification of inactive states – REM and slow-wave sleep, and quiet wakefulness*. Inactive states, i.e. when rats are immobile, were identified as bins having total accelerometer power below the immobility threshold (see ‘Processing of accelerometer data’ above). Slow-wave sleep (SW) states were distinguished by high delta power relative to the delta power threshold (see ‘Processing of the LFP’ above), and similarly, REM sleep states (RM) were distinguished by high theta power relative to the theta threshold (see above) (Eschenko et al., 2006; Gervasoni et al., 2004). The ratio index *delta/(delta + theta)* was then used as a secondary measure to distinguish SW and RM from non-sleep states, with an index > 0.4 indicating SW and an index < 0.4 indicating RM (Eschenko et al., 2006; Gervasoni et al., 2004), such that only epochs with high delta power and ratio index > 0.4 were identified as SW, and only epochs with high theta power and ratio index < 0.4 were identified as RM. The immobile epochs that were not classified as sleeping were labelled as quiet wakefulness. Consistent with prior studies (Gervasoni et al., 2004), we found that rats spent most of their time sleeping (49±6%, n=2) or resting (17±8%) (Figure S5B). Sleep occurred at all hours, but was more frequent when lights were on (51±11%) versus when they were off (38±17%), consistent with the nocturnal habits of rats (Figure S5C).

*Classification of active states – grooming, eating, and active exploration*. Active states were characterized by the accelerometer power being above the immobility threshold.

*Grooming* epochs are characterized by repetitive licking, stroking and scratching. To identify these rhythmic behaviors, we calculated the periodogram of the accelerometer power in 1 s bins and evaluated the power in the 5-50 Hz frequency range. The distribution of these values had a Gaussian shape with a non-Gaussian long tail towards high values. The threshold for high power, indicating rhythmic movements associated with repetitive grooming, was set at the 95^th^ percentile of the Gaussian distribution.

*Eating* epochs were classified based on a strong common-mode oscillatory signal on all electrodes, likely due to electrical interference from mastication. To extract eating epochs, the raw spike data was down-sampled 10-fold (from 30 kHz to 3 kHz), filtered by a 2^nd^ order Butterworth filter and smoothed with a 0.3 s averaging window. We computed the spectrograms for the filtered data in 1 s windows and calculated the total power in the 0-50 Hz frequency band. The distribution of these values had a Gaussian shape with a non-Gaussian long tail towards high values. The threshold for eating was set to the 95^th^ percentile of the Gaussian distribution.

*Task engagement* was identified based on lever-press event times. If less than 2 s had elapsed between presses (corresponding to the average trial length) the time between them was classified as task engagement. Active epochs that were not classified as grooming, eating or task engagement were classified as *active exploration*. The algorithm labeled each time bin exclusively, i.e. only one label was possible for each bin. Bins corresponding to multiple states (e.g. eating and grooming) where classified as *unlabeled* (UL), as were times when no data was available due to brief pauses in the recording.

*Removal of seizure-like epochs*. Long Evans rats are susceptible to absence seizures, characterized by strong ∼8 Hz synchronous oscillations in neural activity and associated whisker twitching (Berke et al., 2004; Gervasoni et al., 2004; Nicolelis et al., 1995). To identify seizure-like episodes, we calculated the autocorrelation of the raw spike data on all electrodes in 2 s windows. To identify peaks associated with seizure-like states, we calculated the periodogram of each autocorrelation function and evaluated the power in the 7-9 Hz frequency range. The distribution of these values had a Gaussian shape with a non-Gaussian long tail towards high values. The threshold for seizure-like states was set to the 95^th^ percentile of the Gaussian distribution. Using this classification, 11% of time-bins were classified as seizure (10 ± 2% for the DLS rat and 12 ± 3% for the MC rat) similar to previously published reports (Shaw, 2003). These epochs were removed from the behavioral state analysis.

*Benchmarking the state classification algorithm*. We benchmarked our automated classification algorithm against a human observer scoring 12 hours of videos of the rats’ behavior. For every state scored manually (superscript ‘m’) we calculated the distribution of the algorithm scores of the same time bins (superscript ‘a’).

|  | Inactive<sup>a</sup> | Grooming<sup>a</sup> | Exploring<sup>a</sup> | Eating<sup>a</sup> | Task<sup>a</sup> | Seizure<sup>a</sup> | Unlabeled<sup>a</sup> |
| --- | --- | --- | --- | --- | --- | --- | --- |
| Inactive<sup>m</sup> | 82.2 | 0 | 0.7 | 0.5 | 0 | 5 | 11.6 |
| Grooming<sup>m</sup> | 1.7 | 29.4 | 43.2 | 0.3 | 6.1 | 1.8 | 17.5 |
| Exploring<sup>m</sup> | 3.1 | 9.4 | 56.3 | 0.5 | 5 | 4.6 | 21.1 |
| Eating<sup>m</sup> | 0 | 1.5 | 17.3 | 79 | 0 | 0 | 2.2 |
| Task<sup>m</sup> | 0 | 1 | 3 | 0 | 96 | 0 | 0 |

While the labels largely overlapped, a significant discrepancy was seen in the identification of ‘grooming’. This is because much of the grooming is not repetitive enough to be distinguished from other exploratory behaviors. However the vast majority of grooming episodes picked out by our algorithm were indeed classified as grooming also by the human observer. Thus the more accurate description of grooming in our analysis would be ‘repetitive’ or ‘periodic’ grooming.

*Calculating state transition times*. We did autocorrelation analysis to estimate mean state transition times, as follows. First, for every state, we constructed a time-varying binary vector indicating whether that state was present (1) or not (0) in 1 s time-bins. We computed the state’s autocorrelogram from this vector over time-lags ranging from 0 to 10000 s. We then summed autocorrelograms over all states and recording days, and scaled them between 0 and 1. Autocorrelograms for both rats were not fit well by a single exponential decay function, but rather by the sum of two exponential functions (time-constants = 91.44 s and 1832.90 s for the DLS rat; 94.09 s and 2012.99 s for the MC rat), indicating that state transitions are the result of a fast and a slow process. The fast timescale process was dominant (fast process / slow process amplitude ratio = 2.7 for the DLS rat and 4.9 for the MC rat), hence we report its time-constant in the Results.

### Data storage and computing infrastructure

*Hardware setup*. Our recordings (64-channels at 30 kHz per channel) generate 1 terabyte (TB) of raw data every 2 days. To cope with these demands, we developed a low-cost and reliable high I/O bandwidth storage solution with a custom lightweight fileserver. Each storage server consists of 24 4TB spinning SATA hard disks connected in parallel to a dual socket Intel server class motherboard via the high bandwidth PCI-E interface. The ZFS file-system (bundled with the open source SmartOS operating system) is used to manage the data in a redundant configuration that allows any two disks in the 24 disk array to simultaneously fail without data loss. Due to the redundancy, each server has 60 TB of usable space that can be read at approximately 16 gigabits per second (Gbps). This high I/O bandwidth is critical for data backup, recovery and integrity verification.

*Distributed computing software setup*. To fully utilize available CPU and I/O resources, we parallelized the data processing (Dean and Ghemawat, 2008). Thread-level parallelization inside a single process is the simplest approach and coordination between threads is orchestrated using memory shared between the threads. However, this approach only works for a single machine and does not scale to a cluster of computers. The typical approach to cluster-level parallelization is to coordinate the multiple parallel computations running both within a machine and across machines by exchanging messages between the concurrently running processes. The ‘map-reduce’ abstraction conceptualizes a computation as having two phases: a ‘map’ phase which first processes small subsets of the data in parallel and a ‘reduce’ phase which then serially aggregates the results of the ‘map’ phase. Since much of our data analysis is I/O limited (like computing statistics of spike waveform amplitudes), we developed a custom distributed computing infrastructure for ‘map-reduce’ like computations for the .NET platform. The major novelty in our framework is that rather than moving the output of the map computation over the network to a central node for performing the reduce computations, it instead moves the reduce computation to the machines containing the output of the map computations in the correct serial order. If the output of the map computation is voluminous compared to the output of each step of the reduce computation then our approach consumes significantly less time and I/O bandwidth. We have used this framework both in a virtual cluster of 5 virtual machines running on each of the aforementioned SmartOS-based storage servers and in Microsoft’s commercial cloud computing platform – Windows Azure.

## Analysis of neural data

### Automated spike sorting algorithm

#### Spike identification

We first extract spike snippets, i.e. the extracellular voltage traces associated with action potentials, automatically from the raw data. A schematic of this process is shown in Figure S1. First, the signal from each channel *s_ch_ (t)* is partitioned into 15 s blocks with an additional 100 ms tacked onto each end of the block to account for edge effects in filtering. Then, for each block of each channel, the raw data is filtered with a 4^th^ order elliptical band-pass filter (cut-off frequencies 300 Hz and 7500 Hz) first in the forward then the reverse direction to preserve accurate spike times and spike waveforms. For each sample in each block, the median across all channels is subtracted from every channel to eliminate common mode noise. Finally, a threshold crossing algorithm is used to detect spikes independently for each tetrode (Figure S1). In our recordings, we use a threshold of 50 μV, which corresponds to about 7 times the median absolute deviation of the signal. After detecting a threshold crossing, we find the sample that corresponds to the local maximum of the event, defined as the maximum of the absolute value across the four channels of a tetrode. A 2 ms (64 sample) snippet of the signal centered on the local maximum is extracted from all channels. Thus, each putative spike in a tetrode recording is characterized by the time of the local maximum and a 256 dimensional vector (64 samples x 4 channels).

Each 15 s block of each tetrode can be processed in parallel. However, since the number of spikes in any given 15 s block is not known in advance, the extracted spike snippets must be serially written to disk. Efficiently utilizing all the cores of the CPUs and simultaneously queuing disk read/write operations asynchronously is essential for keeping this step faster than real-time. In our storage server, the filtering/spike detection step runs 15 times faster than real-time. To extract LFPs, we down-sample the raw data 100-fold (from 30 kHz to 300 Hz) by two applications of a 4^th^ order 5-fold decimating Chebychev filter followed by a single application of a 4^th^ order 4-fold decimating Chebychev filter. After extracting the spike snippets and the LFPs, the raw data is deleted, resulting in a 5-10 fold reduction in storage space requirements.

A typical week-long recording from 16 tetrodes results in over a billion putative spikes. While most putative spikes are low amplitude and inseparable from noise, the spike detection threshold cannot be substantially increased without losing many cleanly isolatable single units. Assigning many billion putative spikes to clusters corresponding to single units in a largely automated fashion is critical for realizing the full potential of continuous 24/7 neural recordings.

#### Automatic Processing Step 1. Local clustering and de-noising

This step of the algorithm converts the sequence of all spike waveforms 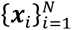 from a tetrode to a sequence of averages of spike waveforms 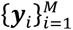 with each averaging done over a set of approximately 100 raw spike waveforms (100*M* ≌ *N*) with very similar shapes that are highly likely to be from the same unit. The output of this step of the algorithm is a partitioning of *N* spike waveforms into *M* groups. 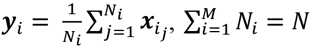 Local clustering followed by averaging produces de-noised estimates of the spike waveform for each unit at each point in time. The goal of this step is not to reliably find all spike waveforms associated with a single unit, but to be reasonably certain that the waveforms being averaged over are similar enough to be from the same single unit. This results in a dataset of averaged spike waveforms that is about a hundred times smaller than the original dataset greatly aiding in speedily running the remaining half of the spike sorting algorithm and in visualizing the dataset.

*Super-paramagnetic clustering*. Clustering is inherently a scale-dependent problem, i.e. the ‘right’ number of clusters in a given dataset depends on the scale being considered. At the coarsest scale, all points can be considered members of a single cluster and at a very fine scale each point belongs to its own cluster. A formal, mathematically precise way of characterizing this tradeoff is to think of clustering as lossy compression (Slonim et al., 2005). The amount of loss is defined as the amount of information lost by replacing each point in a cluster with their ‘average’ and the compression comes from characterizing the entire dataset with just the cluster averages. A simple loss measure is the sum of squared distances between each point in a cluster and the cluster centroid, i.e. the within cluster variance. If each point is assigned its own unique cluster then the loss would be zero. Conversely, if all points were assigned the same cluster then the loss would simply be the variance of the entire set of points. For intermediate amount of loss between these two extremes, the fewest number of clusters, i.e. the largest amount of compression, with at most that much loss, can, in principle, be computed. Conversely for a given number of clusters, one can compute the clustering that results in the smallest amount of loss.

We use the super-paramagnetic clustering (SPC) algorithm (Blatt et al., 1996) in our spike sorting pipeline partly because it parameterizes the loss-compression tradeoff discussed above with a ‘temperature’ parameter. At low temperatures, the algorithm assigns all points to a single cluster. As the temperature parameter is increased new clusters appear until, at very high temperatures, each point is assigned its own cluster. Units with large spike waveforms or very distinctive spike shapes appear at relatively low temperatures. However, other units often appear at relatively high temperatures and clusters at higher temperatures often do not include spikes in the periphery of the cluster. In existing uses of this algorithm for spike sorting the ‘right’ temperature for each cluster is selected manually (Quiroga et al., 2004). Often several clusters at a higher temperature need to be manually merged as they all correspond to the same single unit.

The SPC algorithm also requires a distance measure between pairs of points. In previous approaches to spike sorting, a small number of features (on the order of 10) are extracted from the full 256 dimensional dataset and the Euclidean distance between points in this new feature space is used as the distance measure for clustering (Quiroga et al., 2004). In our experience the number of coefficients that are necessary to adequately capture the distinction between similar, but well isolated units varies substantially depending on the number of units being recorded on a tetrode and the signal-to-noise ratio of the recording. We find that simply using the Euclidean distance in the raw 256 dimensional space avoids this problem without being computationally prohibitive.

*Iterative multi-scale clustering*. Two considerations determine the size of the batch for local clustering. First, SPC requires computing distances between every pair of points, making the algorithm quadratic in the number of points being clustered in one batch. In a typical desktop-class PC, a window size of 10000 spikes runs 8 times slower than real-time, whereas 1000 spike batches run faster than real-time. Second, changes in the spike waveform of a unit over time means that the space occupied by points belonging to a single cluster increases with the size of the temporal window (i.e. batch size), which in turn decreases the separation between clusters. Both of these considerations favor clustering relatively fewer spikes in a batch, and our algorithm does it in batches of 1000 spikes.

However, different neuron types can have very different firing rates (Figure 3). Therefore, a 1000 spike window might contain just a few or even no spikes from low firing rate units. To overcome this problem, we developed a multi-resolution approach for the local clustering step. It identifies clusters corresponding to units with high firing rates, then removes them from the dataset and then re-clusters the remaining spikes, repeating this process iteratively (Figure S2).

*Detailed algorithm for local clustering and de-noising*. The detailed steps of the algorithm for multi-resolution local clustering are described below and a schematic of the whole process is presented in Figure S2.

1. The set of all spike waveforms is partitioned into blocks of 1000 consecutive points. Therefore, the first block contains points {***x***_1_,…, ***x***_1000_}, the second block contains {***x***_1001_,…, ***x***_2000_} and so on.
2. An SPC cluster tree is generated for each block independently. This is computed by first clustering the set of 1000 points at a range of temperatures *T_i_* = 0.01*i*, 0 ≤ *i* ≤ 15. This process assigns a cluster label to each point at each temperature. This matrix of cluster labels is then converted to a tree where each node in the tree corresponds to a cluster at some temperature, i.e. a subset of the 1000 points. The root node of the tree (depth 0) corresponds to a single cluster containing all 1000 points. The children of the root node (depth 1 nodes) correspond to a partition of the set of 1000 points based on the cluster labels at temperature 0.01. For each node in depth 1, the children of that node (depth 2 nodes) correspond to a partition of the points associated with that node based on the cluster labels of those points at temperature 0.02. This is repeated for all temperatures to construct the full tree with depth equal to the number of temperature increments.
3. Each cluster tree is collapsed into a partition (a clustering) of the set of 1000 points (Figure S2B). The simplest technique for getting a partition from an SPC cluster tree is to use all the nodes at a fixed depth, i.e. clustering at a fixed temperature. In practice, this approach suffers from major drawbacks. The lowest temperature at which a cluster first separates from its neighbors varies from unit-to-unit and depends on the signal-to-noise ratio of the spike waveform, how distinct the spike waveform of that unit is, etc. Also, when units appear at relatively high temperatures, the clusters corresponding to single units at those temperatures don’t include many spikes at the periphery of those clusters. Therefore, instead of using a single temperature we recursively merge leaves of the cluster tree based on the loss-compression tradeoff discussed above to generate a partition. This is done by recursively collapsing the cluster tree one level at a time. Specifically,

a. For each leaf node in the cluster tree, the similarity between the leaf node and its parent is first calculated. Let *L* = {*i*_1_, … *i_N_*} be the leaf node which is specified by the indices of the subset of the 1000 points belonging to that node. Similarly, let *P* = {*j*_1_, …, *j_M_*} be the parent of the leaf node. Note that *L* ⊆ *P*. Let 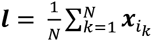 be the average spike waveform of the leaf node and *p* be the average spike waveform of its parent. Let 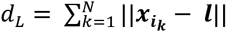 be the total distance of points in the leaf node from their average and 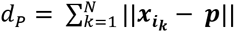 be the distance from the parent node. The difference *d_p_* – *d_L_* = *a_L_* measures how well the parent node approximates the leaf node, i.e. how much additional loss is incurred in approximating the points in the leaf node with the cluster corresponding to the parent node.
b. Let *L* = {*L_i_*} be the set of all N leaf nodes sharing the same parent node *P*. The set of leaf nodes that are poorly approximated by their parent (the well-isolated nodes *I*) are considered distinct clusters. *I* ⊆ *L*, where *L_i_* ∊ *I* if *a_L_i__* > *a*. This encodes the intuition that if a cluster at a given temperature splits into multiple sub-clusters at the next higher temperature that are each quite similar to the parent cluster, then treating each of these sub-clusters as distinct clusters is inappropriate. The parameter *a* provides an intuitive tradeoff between missing distinct clusters that appear at high temperatures and classifying spikes in the periphery of a single cluster into multiple distinct clusters. Let *M* be the number of elements in *I*. If any of the remaining *N* – *M* nodes are well approximated by one of the well-isolated nodes then they are merged together. For *L_i_* ∊ *L/I*, if min_L_j_ *∊I*_ *d_L_j__* – *d_L_i__* < *a*, i.e. if node *L_j_* approximates node *L_i_* well then they are merged. Merging a set of nodes corresponds to creating a node containing all the points from each of the nodes being merged. This yields a set of augmented well-isolated nodes. Any remaining nodes, i.e. non-well-isolated nodes that are also not well-approximated by any of the well isolated nodes, are merged with each other. Therefore, this step results in converting the set of N leaf nodes sharing a parent into a set of *M* or *M* + 1 nodes formed by merging some of them together.
c. A depth *D* tree is converted into a depth *D* – 1 tree by replacing all the leaf nodes and their parents with the merged nodes derived in the previous step.
d. Step a – c are repeated recursively until the tree is of depth 1. The leaf nodes of this tree which are typically vastly fewer in number than the total number of leaf nodes of the original tree correspond to a partition of the set of 1000 points of each block.
4. The centroid of each cluster from the previous step containing at least 15 points contributes one element to the output of the local clustering step, the set of averaged spike waveforms {***y**_i_*} (Figure S2C). In our datasets, clusters with at least 15 spikes in a 1000 spike window correspond to firing rates of approximately 1 Hz or greater, on average. The points belonging to the remaining clusters, i.e. ones with fewer than 15 points, are all pooled together, ordered by their spike time and become the new {***x**_i_*}. The number of spikes in this new subset is approximately 10% of the original. Steps 1-4 are used to locally cluster this new subset of spikes and produce a second set of averaged spike waveforms {***y**_i_*}. This process of re-clustering the low-density clusters is repeated two more times. The averaged spike waveforms from all four scales are then grouped together to form the full set 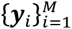 and ordered by the median spike time of set of spikes that were averaged to generate each ***y**_i_*. This process results in an assignment of over 98% of the original set of N spikes to a cluster with at least 15 spikes in one of the 4 scales. The firing rate of units in clusters with at least 15 spikes at the fourth scale is about 0.01Hz in the sample dataset.

#### Automatic Processing Step 2. Sorting and tracking de-noised spike waveforms

This step takes the sequence of averaged spike waveforms 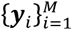 computed in the previous step and identifies the subsets of {***y**_i_*} that correspond to the same putative single unit. A subset of {***y**_i_*} is considered to be the same single unit if distances between ***y**_i_* and ***y**_i_*_+1_ are sufficiently small for the entire subset. As in the previous step, the set 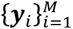 is first partitioned into blocks of 1000 consecutive points and an SPC cluster tree is independently computed for each block, this time for the temperature range *T_i_* = 0.01*i*, 0 ≤ *i* ≤ 10. Then, we use a binary linear programming algorithm inspired by a computer vision problem called segmentation fusion (Vazquez-Reina et al., 2011) to identify the nodes in each cluster tree that correspond to the same single unit.

*Binary linear programming allows units to be tracked over time*. Tracking multiple single units over time from a sequence of cluster trees requires first selecting a subset of the nodes of each cluster tree that correspond to distinct units, followed by matching nodes from adjacent cluster trees that correspond to the same unit. Doing these steps manually is infeasible because of the large volumes of data. In our datasets, the local-clustering step results in ∼100 million averaged spikes from the original set of ∼10 billion spikes per rat. This produces a set of ∼100,000 cluster trees making manual tracking impossibly labor intensive. We instead adapted the segmentation fusion algorithm, invented to solve the problem of reconstructing the full 3D structure of axons, dendrites and soma present in a volume of neuropil from a stack of 2D electron microscopy sections (Kaynig et al., 2015; Vazquez-Reina et al., 2011). A cluster tree is analogous to a multi-resolution segmentation of the 2D image. Identifying nodes across cluster trees that correspond to the same single unit is thus analogous to identifying the same segment across 2D sections as a neurite courses through the neuropil.

The algorithm finds a set of sequences of nodes from the sequence of cluster trees, where each sequence of nodes corresponds to a well isolated single unit. This is done by first enumerating all the nodes in all the cluster trees and all possible links between nodes in adjacent cluster trees. Then a constrained optimization problem is solved to find a subset of nodes and links that maximize a score that depends on the similarity between nodes represented by a link and the ‘quality’ of a node. This maximization is constrained to disallow assigning the same node to multiple single units and to ensure that if a link is selected in the final solution then so are the nodes on each side of the link.

*Details of the binary linear programming algorithm*. Each step of the binary linear programming algorithm is detailed below (see also Figure S3).

1. The sequence of cluster trees is grouped into blocks of 10 consecutive trees with an overlap of 5 cluster trees. Solving the binary linear program for blocks larger than 10 trees is computationally prohibitive.
2. The segmentation fusion algorithm is run independently for each block of 10 cluster trees (Figure S3). Let 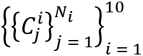 be the set of binary indicator variables representing all nodes in all 10 cluster trees where the cluster tree indexed by *i* contains a total of *N_i_* nodes. The total number of nodes is *N* = 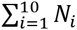. Let 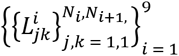 be the variables representing the set of all links between adjacent cluster trees. Link 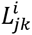 connects clusters 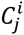 and 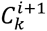. The total number of links is 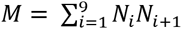. Solving the linear program requires choosing a {0,1} value for each of the *N* + *M* binary variables that maximizes the objective function 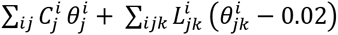. The objective function is a weighted linear sum of all the binary variables where the cluster weights 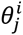 represent the ‘quality’ of the cluster and the link weights 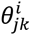 represent the similarity of the clusters joined by the link. The link weights are numbers in the range (0,1). The threshold of 0.02 serves to give negative weight to links between sufficiently dissimilar clusters, effectively constraining the value of the variables representing those links to 0. This objective function is to be optimized subject to three sets of constraints. The first, 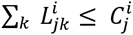, enforces the constraint that if the node variable 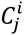 is assigned a value of 1 then out of all the outgoing links from the node 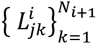, at most one is chosen (Figure S3C). Similarly, the second set of constraints, 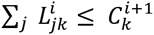, enforces the requirement that at most one incoming link to a node is chosen (Figure S3B). The third set of constraints enforces the requirement that for each of the 1000 points in a cluster tree at most one of the nodes containing that point is chosen (Figure S3D). This translates to inequalities 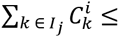 1 where the set of indices *I_j_* represents nodes in the path from the root of cluster tree *i* to the *j^th^* leaf node of the cluster tree. Therefore, the total number of constraints of this type for each cluster tree is the number of leaf nodes in that cluster tree. The link weight 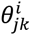 is the Euclidean distance between the average spike waveform of clusters 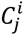 and 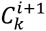 non-linearly scaled by a sigmoid function to fall in the range (0,1). If *d* is the distance then 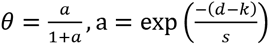, s = 0.005,k = 0.03. The parameter *s* controls the steepness of the sigmoid and the parameter *k* sets the distance *d* at which *θ* = 0.5. The cluster weight 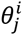 gives preference to clean well-isolated clusters, i.e. clusters that appear at low temperatures and retain most of their points across a large temperature range. Let *N*^(0)^ be the number of points in the cluster corresponding to 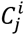. Let 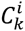 be the largest cluster amongst the child nodes of 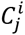 and let *N*^(1)^ be the number of points in 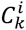. Similarly let *N*^(2)^ be the number of points in the largest cluster among the child nodes of 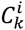. Given the sequence of cluster sizes *N*^(0)^, *N*^(1)^,…, *N^(a)^* where *N^(a)^* is the number of points in a leaf node of cluster tree, 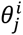 is defined as *N*^(0)^/(*N^(0)^* + … + *N^(a)^*). This measure of cluster quality penalizes clusters that split into smaller clusters at higher temperatures and clusters that only appear at high temperatures.
3. The results of the previous step, i.e. the subset of the M links of each block that maximizes the objective function, are finally combined to produce a sorting that tracks single units over long time periods despite gradually changing waveforms. Links that are part of two instances of the segmentation fusion procedure due to the overlap mentioned in step 1 are only included in the final solution if both linear programs include them. The set of links chosen by the segmentation fusion algorithm are chained together to get long chains of clusters. For instance if links 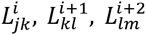 are assigned values of 1 in the solution to the segmentation fusion linear program then all points in clusters 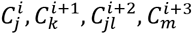 belong to the same chain and hence the same single unit. Each point that does not belong to any chain is assigned to the chain containing points most similar to it (as measured using the sigmoidal distance of step 2) as long as the similarity *θ* >0.02 (again the same threshold as used in step 2).

*Merging the chain of nodes and links identified by the binary linear programming algorithm*. Often, spike waveforms have multiple equal amplitude local extrema. Since the waveforms are aligned to the local extrema with the largest amplitude during the spike identification phase, different waveforms from the same unit can be aligned to different features of the waveform. This results in multiple chains for the same single unit since the Euclidean distance between waveforms aligned to different features is very large. This is remedied by merging chains that contain spikes from the same recording interval if the translation-invariant distance between the spike waveforms of the chains is sufficiently low in the overlap region. The translation invariant distance is computed by first calculating the distance between a pair of spike waveforms for a range of relative shifts between the pair and then taking the minimum of this set of distances. Overlapping chains with the smallest translation-invariant distance are first merged. This is done recursively until either no overlap between chains remains or overlapping chains have distinct spike waveforms and hence correspond to different simultaneously recorded single units.

#### Visualization and manual verification

The final output of the unsupervised spike sorting algorithm consisted of long chains (median length = 8.5 hours) corresponding to single-unit spikes linked over time. However, our algorithm did not link spike chains over discontinuities in recording time (i.e. across recording files), or in the case of rapid changes in spike shape that occasionally occurred during the recording, or when spike amplitudes drifted under 75 μV for brief periods. In such cases, we had to link, or ‘merge’ chains across these discontinuities.

In order to visualize, merge and manually inspect the unsupervised chains, we developed a MATLAB program with a graphical user interface (GUI). This allows users to semi-automatically merge chains belonging to the same unit across discontinuities based on end-to-beginning similarity in their spike waveforms and inter-spike interval distributions. We perform these merging events only if the time-gap between the end of one chain and the start of the next is less than 5 hours of recording time, or up-to 24 hours in the case of gaps in recording time (which in the two experiments we report on were, on average, 2 hours). In occasional cases of inaccurate spike sorting by the unsupervised algorithm, either when spike amplitude was too close to noise or when chains’ spike waveforms were too similar, we manually split the clusters using MClust (MClust 4.3, A. D. Redish et al).

*Cluster quality*. After spike sorting (see below and Supplementary Experimental Procedures), we computed standard measures of cluster quality, specifically the isolation distance (Harris et al., 2000), L-ratio (Schmitzer-Torbert et al., 2005) and fraction of inter-spike intervals less than 1 ms, for every unit within consecutive one-hour blocks. Isolation distances and L-ratios were calculated with four features for every electrode channel – the waveform peak, waveform energy (L^2^ norm), and the projections of the spike waveforms on the first two principal components of all spike waveforms in the one-hour block. Only units meeting all of our quality criteria (isolation distance => 25, L-ratio <= 0.3, fraction ISI less than 1 ms <= 0.01) in a particular block were included in further analysis.

*Comparison to manual spike sorting*. We performed manual spike sorting for 2 separate hour-long blocks from a continuous electrophysiology dataset recorded in the dorsolateral striatum. Manual sorting was performed with M-Clust 3.5 (A. D. Redish et al). The resultant clusters were then compared to those obtained by our unsupervised algorithm as described in the Results and Figure S4A-C.

*Discontinuous tracking of units across days*. We identified units present (>= 50 spikes) in daily one-hour ‘sessions’ from 10 to 11 am, during execution of a skilled motor sequence task (Kawai et al., 2015). We computed four measures of unit similarity across consecutive days’ sessions (Figure S4D), namely the Euclidean distance and correlation coefficient (Emondi et al., 2004) between average unit spike waveforms (waveforms recorded on different channels of a tetrode were concatenated together to yield a 256-element vector), the correlation coefficient between units’ inter-spike interval histograms (Dickey et al., 2009) (see below), and the difference in firing rates (Fraser and Schwartz, 2012). Correlation coefficients were Fisher transformed, and spike waveform Euclidean distance and firing rates were log-transformed to appear more Gaussian (Dickey et al., 2009; Fraser and Schwartz, 2012) (Figure S4D). Similarity metrics were calculated for all pairs of units recorded on the same tetrode across consecutive sessions/days and classified as either the ‘same’ or ‘different’ as per our FAST algorithm. Linear discriminant analysis (LDA) (Tolias et al., 2007) was then used to identify a similarity threshold that would best separate ‘same’ from ‘different’ unit pairs at false positive error rates ranging from 25% (Dickey et al., 2009) to 5% (Fraser and Schwartz, 2012), 1% and 0.25%. Here, false positive rates are defined as the proportion of ‘different’ units misclassified by the classifier as the ‘same’. We also tried a quadratic classifier (Dickey et al., 2009; Fraser and Schwartz, 2012) but found no significant difference in performance compared to the linear classifier. LDA fits multidimensional Gaussians to the similarity distributions identified by ‘same’ or ‘different’ class labels. Therefore, assuming a uniform prior, we can calculate the posterior probability of a unit pair being part of the ‘same’ distribution and use it as a similarity score. Units were matched across consecutive daily ‘sessions’ if their similarity scores exceeded the threshold set by the expected false positive rate, with highest similarity unit pairs being matched first (Emondi et al., 2004).

*Identification of putative cell types*. We used mean spike waveforms and average firing rate (Barthó et al., 2004; Berke et al., 2004) to separate units into putative cell types: medium spiny neurons (MSNs) and fast spiking neurons (FS) in the striatum, and regular spiking (RS) and fast spiking (FS) neurons in the cortex. Briefly, we performed kmeans clustering of units based on their peak-normalized spike-waveform, concatenated with their log-transformed firing rates. The number of clusters (k=2) was chosen by visual inspection of unit waveforms in three dimensions - the first and second principal components of the spike waveform, and unit firing rate.

*Firing rate stability*. Firing rate similarity compared across different days *i* and *j* for the same unit, or when comparing distinct units *i* and *j* recorded on the same day, was measured by the following formula:

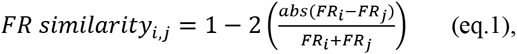

where *FR* is the firing rate. A firing rate similarity score of 1 means that the two firing rates *FR_i_* and *FR_j_* are identical while a firing rate similarity score of -1 implies maximum dissimilarity in firing rates, such that one of the firing rates is 0. When comparing firing rates for the *same unit across time*, we calculated average firing rate similarity scores for time-lags ranging from 1 to 10 days. When comparing firing rates *across units* and within the same time-bin, we averaged together all similarity scores for pairs of units of the same putative unit-type recorded in that time-bin.

*Stability of inter-spike interval (ISI) distribution*. We computed ISI histograms using 50 log-spaced time-bins spanning inter-spike intervals from 1 ms to 1000 s. We used Pearson’s correlation coefficient to measure the similarity between ISI distributions on different days for the same unit, and between ISI distributions of distinct units recorded on the same day. For each unit, as with the firing rate similarity measure described above, we averaged the ISI correlations for each time-lag ranging from 1 to 10 days. When comparing across units, we only compared their ISI distributions to other simultaneously recorded units of the same putative unit-type.

*Stability of state-tuning*. We used the Pearson’s correlation coefficient to estimate the similarity between units’ state-dependent firing rate profiles (i.e. their state-tuning), either across days, or between different units recorded simultaneously.

*Spike-triggered averages (STA)*. The spike-triggered average (STA) is commonly used in sensory neuroscience to characterize a neuron’s ‘receptive field’ in stimulus space (Marmarelis and Naka, 1972; Meister et al., 1994). We adapted this method to estimate ‘response fields’ in movement space (Cheney and Fetz, 1984; Cullen et al., 1993; Serruya et al., 2002; Wessberg et al., 2000). For each unit, we computed spike-triggered averages of accelerometer power in three different behavioral states – eating, grooming and active exploration. The accelerometer power *P_acc_* was calculated as:

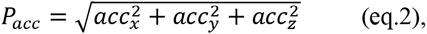

where *acc_i_* is the accelerometer signal on channel *i*. The accelerometer power was band-pass filtered between 0.5 Hz and 300 Hz using a 3^rd^ order Butterworth filter and zero-phase filtering. Spike-triggered averages (STA) were computed for each unit in a particular behavioral state by averaging, over spikes, the accelerometer power in a window of ±500 ms centered on each spike. Formally,

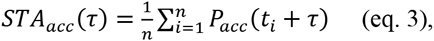

where τ is the time-lag, sampled at 3.33 ms resolution, between spike and accelerometer power ranging from -500 to +500 ms, and *t_i_* the arrival times of unit spikes (a total of *n* spikes) in a particular behavioral state. We only considered ‘trigger-spikes’ that were at least 500 ms away from a behavioral state-transition.

To measure the significance of the resultant STA kernels, we computed noise STAs after jittering spike times by ±500 ms. We used the standard deviation of the noise STAs across time-bins to calculate the Z-scored STAs. Any STA having at least 2 bins equal to or exceeding a Z-score of 5 was considered statistically significant and used for further analysis.

Just as with ISI distribution similarity, we calculated the similarity between STAs for the same unit on different days using Pearson’s correlation coefficient. We also compared similarity of STAs across simultaneously recorded units, but only within the same behavioral state.

*Peri-event time histograms (PETHs)*. We computed peri-event time histograms (PETHs) of instantaneous firing rates aligned to specific behavioral events during execution of a lever-pressing task (Kawai et al., 2015). We computed PETHs in windows ±200 ms around the first lever-press event of a trial, as well as at times of nose-pokes into the reward port following successful trials. In order to restrict our analysis of neural activity to periods of stereotyped behavior, we selected only rewarded trials that followed previously rewarded trials (to control for starting position), and these trials’ inter-press intervals had to be within 20% of the target inter-press interval of 700 ms (to ensure movement stereotypy). To estimate a unit’s instantaneous firing rate on each trial, we convolved its event-aligned spike train with a Gaussian kernel (σ = 20 ms). These firing rates were then averaged over the selected trials from a given day to yield the unit PETH. To determine whether a particular PETH had statistically significant modulation in firing rate, we estimated the p-value and Z-score for each bin in the PETH using a bootstrap approach. P-values were then pooled across days for all lever-press or nose-poke PETHs for a particular unit using Fisher’s method. A unit was deemed to have significant modulation in its time-averaged PETH if at least 2 bins had p-values < 1e-5. At this significance threshold the probability of getting > 1 false positive over 1000 neurons and 20 time-bins is less than 0.01.

Similarity across PETHs for the same unit across days, or between different units on the same day was computed using the Pearson’s correlation coefficient, similar to the measurement of similarity between ISI distributions and STAs. Similarities in lever-press and nose-poke PETH similarities were averaged for each unit for a particular time-lag (ranging from 1 to 10 days).

*Cross-correlograms*. We computed cross-correlograms for all unit pairs recorded simultaneously using spikes recorded in all labeled behavioral states (Figure 4) or in specific behavioral states (Figure S6). For each unit pair, we only calculated correlograms on days on which both units were recorded for at least 60% of the time (i.e., >14 of 24 hours). Correlograms were calculated in 1 ms bins and smoothed with a 3 ms boxcar filter. Correlograms for unit pairs recorded on the same tetrode are, by design, zero at zero lag since only one spike event is allowed per tetrode in a 1 ms bin. For these pairs, the average of the preceding and following bins was used to approximate the value of the correlogram at zero lag. Correlations were considered significant if two consecutive bins of the correlogram were above a threshold set at 3 standard deviations of a shuffled correlogram computed for the same pair (one spike train was shuffled by adding ±400 ms random jitter to each spike). Only correlograms that had at least 5000 spikes could be considered significant. Each significant correlogram was characterized by its maximum peak (positive correlation) or trough (negative correlation), averaged over 3 bins around the peak/trough and normalized by the average bin height of the shuffled correlogram. Thus a peak > 1 indicates a positive correlation whereas a trough < 1 indicates negative correlations. The lag was defined as the bin location of the peak/trough in the correlogram.

*Stability of correlograms*. We used the Pearson’s correlation coefficient to quantify the similarity between correlogram distributions on different days, or in different states for the same unit pairs.

*Firing rate correlations*. We first calculated each unit’s instantaneous firing rate, smoothed at different timescales by convolving its spike-train with a Gaussian kernel with a standard deviation of 0.1, 1, 10, 100 or 900 s. For each unit pair, we computed Pearson’s correlation coefficient between their instantaneous firing rates recorded over the course of a day, and then averaged these correlation coefficients calculated over all days on which the unit pair was simultaneously recorded. For each unit, we only considered days on which a unit was recorded for at least 60% of the time (>14 of 24 hours). For firing rate correlations computed during slow-wave sleep alone, we only considered time-bins during which rats were in slow-wave sleep at least 80% of the time.

*Tests of statistical significance*. We used the Kruskal-Wallis one-way analysis of variance test to determine whether differences in within-unit similarity measures of firing rate, ISI distribution, interneuronal correlograms, behavioral state modulation of firing rate, spike-triggered averages and PETHs, over time-lags of 1 to 10 days were significant when compared to each other or to similarity measures computed across units. P-values were corrected for multiple comparisons by the Tukey-Kramer method. We used an alpha value of 0.05.

